# C-STEM: ENGINEERING NICHE-LIKE MICRO-COMPARTMENTS FOR OPTIMAL AND SCALE-INDEPENDENT EXPANSION OF HUMAN PLURIPOTENT STEM CELLS IN BIOREACTORS

**DOI:** 10.1101/2021.07.05.451086

**Authors:** Philippe J.R. Cohen, Elisa Luquet, Justine Pletenka, Andrea Leonard, Elise Warter, Basile Gurchenkov, Jessica Carrere, Clément Rieu, Jerome Hardouin, Fabien Moncaubeig, Michael Lanero, Eddy Quelennec, Helene Wurtz, Emilie Jamet, Maelle Demarco, Celine Banal, Paul Van Liedekerke, Pierre Nassoy, Maxime Feyeux, Nathalie Lefort, Kevin Alessandri

## Abstract

Human pluripotent stem cells (hPSCs) have emerged as the most promising cellular source for cell therapies. To overcome scale up limitations of classical 2D culture systems, suspension cultures have been developed to meet the need of large-scale culture in regenerative medicine. Despite constant improvements, current protocols relying on the generation of micro-carriers or cell aggregates only achieve moderate amplification performance. Here, guided by reports showing that hPSCs can self-organize *in vitro* into cysts reminiscent of the epiblast stage in embryo development, we developed a physio-mimetic approach for hPSC culture. We engineered stem cell niche microenvironments inside microfluidics-assisted core-shell microcapsules. We demonstrate that lumenized three-dimensional colonies maximize viability and expansion rates while maintaining pluripotency. By optimizing capsule size and culture conditions, we scale-up this method to industrial scale stirred tank bioreactors and achieve an unprecedented hPSC amplification rate of 282-fold in 6.5 days.

**TEASER:** Optimizing human pluripotent stem cells amplification by recapitulating and protecting biomimetic colonies in a bioreactor

## INTRODUCTION

Following pioneering works targeting Parkinson’s disease (*1, 2*), diabetes (*3, 4*), macular degeneration (*5*) or heart failure (*6, 7*), cell therapies are now addressing an increasing diversity of indications (*8, 9*). Therapeutic cells represent a hope for millions of patients worldwide with chronic diseases or unmet medical needs. For each of those patients, the number of required cells range between 10^5^ and 10^10^ cells. While academic pre-clinical studies or small-scale clinical trials have already been proved to be successful, a transition to true clinical scale is now urgently needed. To enable treatment of thousands to millions of patients, production capacity must scale while maintaining high-quality cells compatible with transplantation.

Due to their unlimited self-renewal capacity and potency to give rise to any cell type in the body, human pluripotent stem cells (hPSCs) hold great promise to provide the required quantities of therapeutic cells. *In vivo*, a few embryonic stem cells can give rise to the 30×10^12^ cells of the adult human body (*10*) while maintaining their genome integrity. As a consequence, much effort has been dedicated to the isolation and *in vitro* culture of these cells in physiological conditions.

Historically, the first stem cell lines were established by micro-dissecting embryos and manual passaging as two-dimensional (2D) adherent epithelial colonies on a layer of feeder cells (*11–13*). Cell proliferation and pluripotency maintenance were thus achieved *in vitro*. The need for an embryonic source was then alleviated with the discovery that differentiated cells could be reprogrammed into induced pluripotent stem cells (iPSC) (*14*). This breakthrough led to pioneer works in developmental cell biology and catalyzed the development of cell therapy applications. Even though a lot of work has been devoted to the development of media and substrate compatible with clinical use, topology of hiPSCs cultures has not been gathering the same attention. Nevertheless, 2D *in vitro* cultures suffer high mortality rates, spontaneous differentiation and genetic drift (*15*). An estimated 40-fold increase in the number of mutations compared to *in vivo* conditions (*16*) often results in rejection of clinical batches due to safety concerns (*17, 18*). In addition, 2D culture systems have limited “scale-out” possibilities: for example, generating 1 trillion pluripotent stem cells, would require almost 1000 m^2^ of plastic dishes and countless handmade passaging (*15*). The non-scalability and the inability to conserve high quality cells remain the main limitations of 2D culture systems for clinical applications (*18*).

More recently, 3D hPSCs cultures have been developed to model developmental processes. After seeding stem cells in bulk extracellular matrix (ECM) (*19, 20*) or using microfluidic chips (*21*), cell clusters self-organize into a monolayered epithelium recapitulating numerous features of an epiblast that generates *in vivo* all tissues in an amniote embryo through gastrulation. Even though these elegant approaches allowed to gain much insight into morphogenesis mechanisms (*22*), they are not designed for the production of hPSCs per se.

From a bioproduction perspective, significant advances have been achieved by developing 3D suspension cultures and bioreactors. Whether hPSCs are grown as floating aggregates or adherent at the surface of microcarriers, those systems provide increased surface-to-volume ratio and scale-up potentialities. Additionally, the use of stirred-tank bioreactors (STBR) allowed better control of the culture parameters without direct human intervention. Indeed, stirring-mediated mechanical agitation avoids sedimentation of the aggregates or microcarriers onto the bottom of the culture vessel and heterogeneities in the culture medium compositions. Nonetheless, one intrinsic drawback of agitation is that hydrodynamic shear induces cellular damages (*23*). This effect is all the more dramatic as the vessel increases in size: the larger the volume of the bioreactor is, the more power per unit of volume needs to be injected for effective homogenization. Even though expansion rates with hPSC aggregates are regularly increased and could reach up to 70-fold within 7 days (*24, 25*), the ability of scaling up the volume of the production is still restricted. Typically, this mechanical upper limit makes it difficult to reach batches larger than 1 liter or equivalently few billion cells. Finally, the current bottleneck for industrial scalability is the difficulty to fulfill simultaneously high amplification rate, volume and physiological quality.

In this work, by bridging the gap between two distinct communities, namely developmental biology and stem cells bioprocessing, we show that recapitulation of hPSC niche-like micro-environments allows to optimize the suspension culture of hPSC in STBRs. More precisely, we propose a system that utilizes a high throughput microfluidic encapsulation technology compatible with suspension culture of stem cells in a bioreactor and amenable to the production of large volume batches without any degradation of cell survival in contrast with 2D colony-based systems for instance. Briefly, iPSCs are encapsulated in alginate hollow microcapsules internally coated with ECM components at low cell seeding concentration. We assess the maintenance of stemness and pluripotency upon 3D culture in suspension. We then characterize the cell growth inside the capsules before showing that upscaling to 10 liter stirred-tank bioreactor allows to reach unrivalled amplification factors. Altogether, the C-STEM technology overcomes the scale-up bottleneck faced in cell therapy bioproduction. We discuss that the origins of this performance could be related to unprecedented cell viability of 3D lumenized colonies within these capsules.

## RESULTS

### High-throughput microfluidic encapsulation of hiPSCs

Using a microfluidic technique to generate hollow hydrogel spheres (Fig. 1A, Movie S1, and detailed description in the Materials and Methods section) previously developed by us and others (*26–29*) we encapsulated human induced pluripotent stem cells (hiPSC) in liquid core capsules. Briefly, the working principle is the following: a three-layered cylindrical flow is generated using a co-extrusion microfluidic device: the cell suspension is surrounded by an alginate solution. Both solutions are separated by a sorbitol solution to prevent diffusion of calcium released from the cell suspension towards the alginate solution. Upon exiting the microfluidic device at high flow rate (on the order of 120 ml/h for all three solutions), the liquid jet is fragmented into small droplets due to the Plateau-Rayleigh instability. By contrast with the dripping regime obtained at lower flow rate that gives rise to droplets of size in the order of the capillary length (i.e. ∼mm), the droplet radius is here dictated by the extruder’s nozzle size (∼200 microns diameter) (*30*). This reduced size, which is below the distance over which oxygen and nutrients supply is limited within a tissue, allows to avoid the formation of a necrotic core (*31*). Once they fall in a calcium bath, the alginate solution droplets undergo gelation and trap the cell suspension in their interior. Our routine protocol produces capsules at a rate of about 3 kHz (Fig. S1), meaning that a 30 seconds operation generates 100,000 capsules. Morphological analysis shows that capsules are monodisperse in size with mean external radius R = 205 µm±39 µm. and that their shape is close to spherical, with a circularity parameter C= 0.84 ± 0.04 (Fig. 1B).

**Fig. 1.**
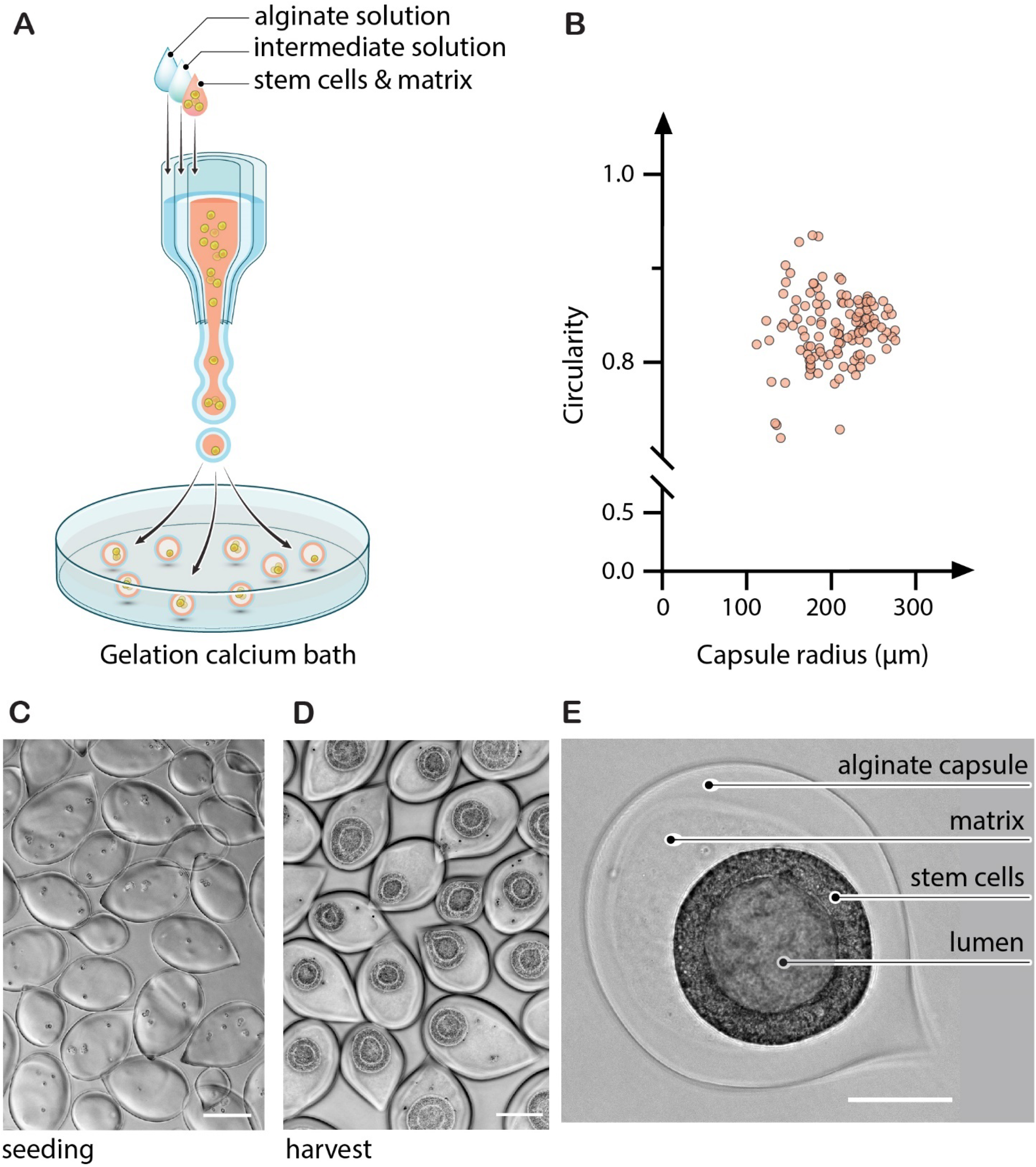
Encapsulation of human pluripotent stem cells (hPSCs) and suspension culture of 3D lumenized colonies. (**A**) Working principle of the microfluidic encapsulation technique. Co-extrusion of three co-axial flows generates composite cell-and extracellular matrix (ECM)-laden droplets. The outer layer composed of alginate solution undergoes gelation upon contact with a calcium bath. Cells are entrapped in the core-shell capsules and ECM condensates onto the internal wall to form a niche-like environment. (**B**) Morphometric analysis of the capsules. Graph of capsule circularity as a function of capsule radius for a representative batch of capsules (n=125). (**C-D**) Phase contrast micrographs of the encapsulated hPSCs after seeding at day 0 (C) and before harvest at day 7 (D) of the suspension culture course. Scale bar is 200µm. (**E**) Magnified phase contrast image showing the hollow alginate capsule revealing the cyst architecture of the encapsulated hPSC colony. Scale bar=100 µm.

In order to provide a niche-like environment to hiPSCs, Matrigel, an ECM mixture, is co-injected with the cell suspension (*28*). Empirically, we found that a minimal volume fraction of 25% was required to form a continuous matrix layer anchored to the inner wall of the capsule, with the excess (if any) being found as floating gel pieces inside the capsule (*29*). Most of the experiments reported in these works were performed with 50% of Matrigel in volume fraction. The granularity seen in the core of the capsule (Fig. 1E) thus corresponds to small floating aggregates of ECM. Most encapsulations reported hereafter were performed with a density of 0.4×10^6^ cells/ml in the cell/matrix suspension, unless otherwise stated, and led to a mean number of cells per capsule right after encapsulation (day 0) of ∼2.5 (Fig. 1C), meaning that ∼10% of the capsules are empty, consistently with a Poisson distribution. After 6 to7 days of culture, practically defined as the harvest time under these seeding conditions, 3D colonies of hiPSCs were observed, suggesting, not only that hiPSCs survive, but also that they could proliferate (Fig. 1D). Higher magnification reveals the presence of a lumen (Fig. 1E).

### Culture, expansion and stemness of encapsulated 3D hiPSC colonies

Using phase contrast imaging, we observed in more details the growth kinetics of these 3D hiPSC colonies. First, hiPSCs form a small cluster (typically during the first 24h) before self-organizing in a cyst structure around a central lumen (Fig. 2A, Movie S2 & S3). Then, the 3D hiPSC colonies grow within the capsules while keeping the same spherical shape. In the early stages, the cells within the monolayer of the cyst have a cuboid cell shape of about 10 µm side (Fig. 2B left, ∼5 days post-encapsulation). Before harvesting, as seen in the confocal image of a representative 7 day-old hiPSC 3D colony immunostained for actin and nucleus, cells exhibit an elongated morphology perpendicular to the surface of the cyst. Yet, the cyst remains monolayered suggesting a transition towards a pseudostratified columnar epithelium with most nuclei being located on the basal side opposing the lumen. In this stage, the cysts are characterized by a thickness of about ∼40 µm (Fig. 2B right) and a radius of about ∼100 µm Fig. S2, day 7). Note that later stages are ignored. Indeed, if cells are not harvested, cysts become confined by the capsules and further grow inwards leading to a progressive loss of the lumen (Movie S4) and eventually the appearance of “fractures” (Fig S2).

**Fig. 2.**
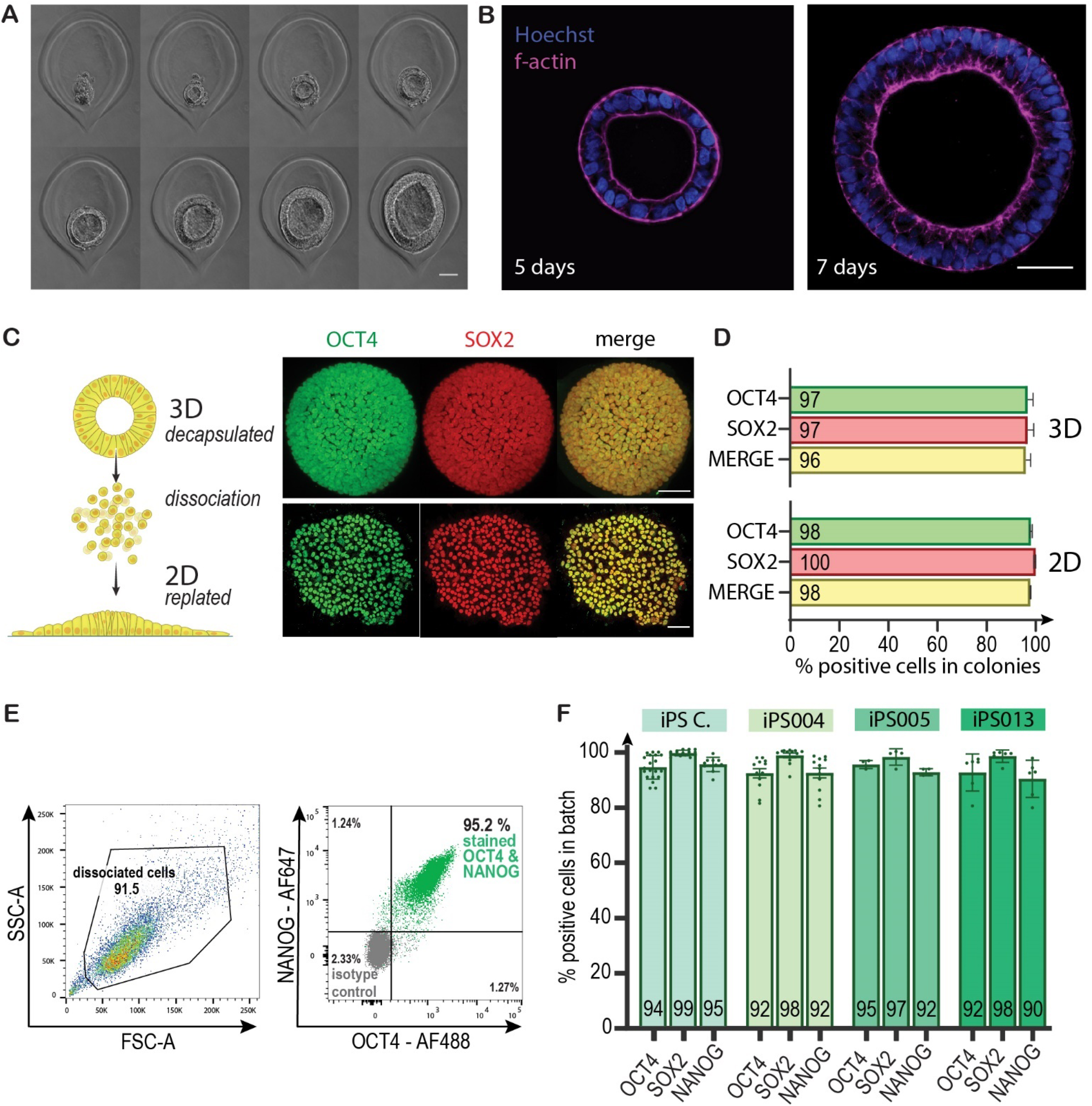
Morphology, growth and stemness of *in capsulo* self-assembled 3D hPSC colonies. (**A**) Snapshots of phase-contrast microscopy images showing the growth of an 3D hPSC colony. The time interval between successive images is 12h. The scale bar is 50µm. (**B**) Confocal image of the equatorial plane of a hPSC colony grown in a capsule at day 5 (left) and day 7 (right)cytoskeletal actin is stained in purple (Phalloidin) and nuclei in blue (Hoechst). Scale bar=50µm. (**C**) Left: Cartoon explaining how 3D hPSC colonies are dissociated and replated to form 2D colonies. Right: Immunostaining of a representative encapsulated 3D colony (upper panel, scale bar=50µm) and a 2D colony obtained after replating (lower panel, scale bar=100µm): OCT4 (green) SOX2 (red) and Merge (OCT4/SOX2). (**D**) Percentage of cells positive for markers of stemness among 4 representative colonies co-stained for OCT4 and SOX2 in 3D capsules (upper panel) and in 2D (lower panel). Number of counted nuclei: n = 1159 for 3D and n=671 for 2D cells, see also Fig. S3). Error bars show the standard deviation of the mean. (**E**) Flow cytometry dot-plots for stemness markers (OCT4 and NANOG) of a batch of 3D hPSC colonies after 7 days of culture (T-Flask). (**F**) Histograms showing the percentage of OCT4, SOX2 and NANOG positive cells in 3D hPSC colonies (culture in T-Flask) analyzed by flow cytometry at 7 days post encapsulation for 4 iPS cell lines (with n>=3 independent biological replicates per cell line, n=42 total number of experiments). Error bars represent the standard error of the mean.

The maintenance of the stemness of the encapsulated 3D hiPSC colonies was then checked. The expression of key self-renewal markers such as OCT4 and SOX2 was first assessed after capsule dissolution, fixation and staining. The alginate shell was dissolved by adapting the protocol of (*29*): we performed a short rinse of ReLeSR, which serves here as a calcium chelator that gently dissolves the alginate gel. Fixation and immunostaining were carried out following standard protocols (see Materials and Methods section) while preserving the 3D architecture. Image analysis of representative confocal images of individual cysts (Fig. 2C (top row) and Fig. S2) allows to derive that the percentage of cells positive for OCT4 and SOX2 is 97% (Fig. 2D top row). To further assess the consistency of stemness phenotype, we applied the approach pursued in a different context for epiblast-stage hPSCs spheroids by Freedman et al. (*32*). “Naked” hiPSCs cysts were dissociated and replated into 2D cultures (Fig. 2C). We observed that 2D colonies are readily formed and stemness markers are detected (Fig. 2C-D, bottom row) with a percentage of OCT4 ansd SOX2 positive cells larger than 98%.

Following this characterization at the scale of individual capsules, we sought to assess the potential variability between capsules and between hiPSC lines. We thus dissociated the bulk suspension cultures, extended staining to OCT4, SOX2 and NANOG and performed flow cytometry (Fig. 2E-F). For 4 different cell lines (see Materials and Methods section) by pooling all experiments (n = 42) of each hiPSC line, we found that the mean percentage of positive cells is 93% OCT4, 98% SOX2 and 92% NANOG (Fig. 2F). This finding is in good agreement with the above-described findings at the single 3D colony level, suggesting an overall homogeneity of the stem cell culture.

Note that, if capsules culture was prolonged beyond 7 days, despite drastic changes of topology from cyst to aggregate (Fig. S2 C-E and Movie S4), the stemness of the hiPSCs was not affected, as revealed by OCT4 and SOX2 staining before and after lumen collapse (Fig. S2 B-C), suggesting that harvest timing is not critically stringent with respect to the stemness maintenance.

### Pluripotency and genomic integrity of encapsulated 3D hiPSC colonies

While OCT4, SOX2 and NANOG are often considred as pluripotency markers, they actually are stemness markers. To assess more thoroughly the pluripotency and validate the quality of hPSCs upon 3D culture in their ability to differentiate as bona fide pluripotent stem cells, the most commonly used assay is the trilineage differentiation assay (*33*), in which stochastic differentiation is induced. Following a standard protocol (*34*) (see Materials and Methods section), decapsulated and dissociated hiPSC (from the three available cell lines) were driven towards early differentiation, as shown by the stainings for specific markers of the 3 germ layers: namely b3-tubulin (TUJI) for ectoderm, smooth muscle actin (α-SMA) for mesoderm and a-fetoprotein (AFP) for endoderm (Fig. 3A). Even though the expression level may differ from one cell line to another, all stainings are positive and clearly reveal a differenciation into the three germ layers.

**Fig. 3.**
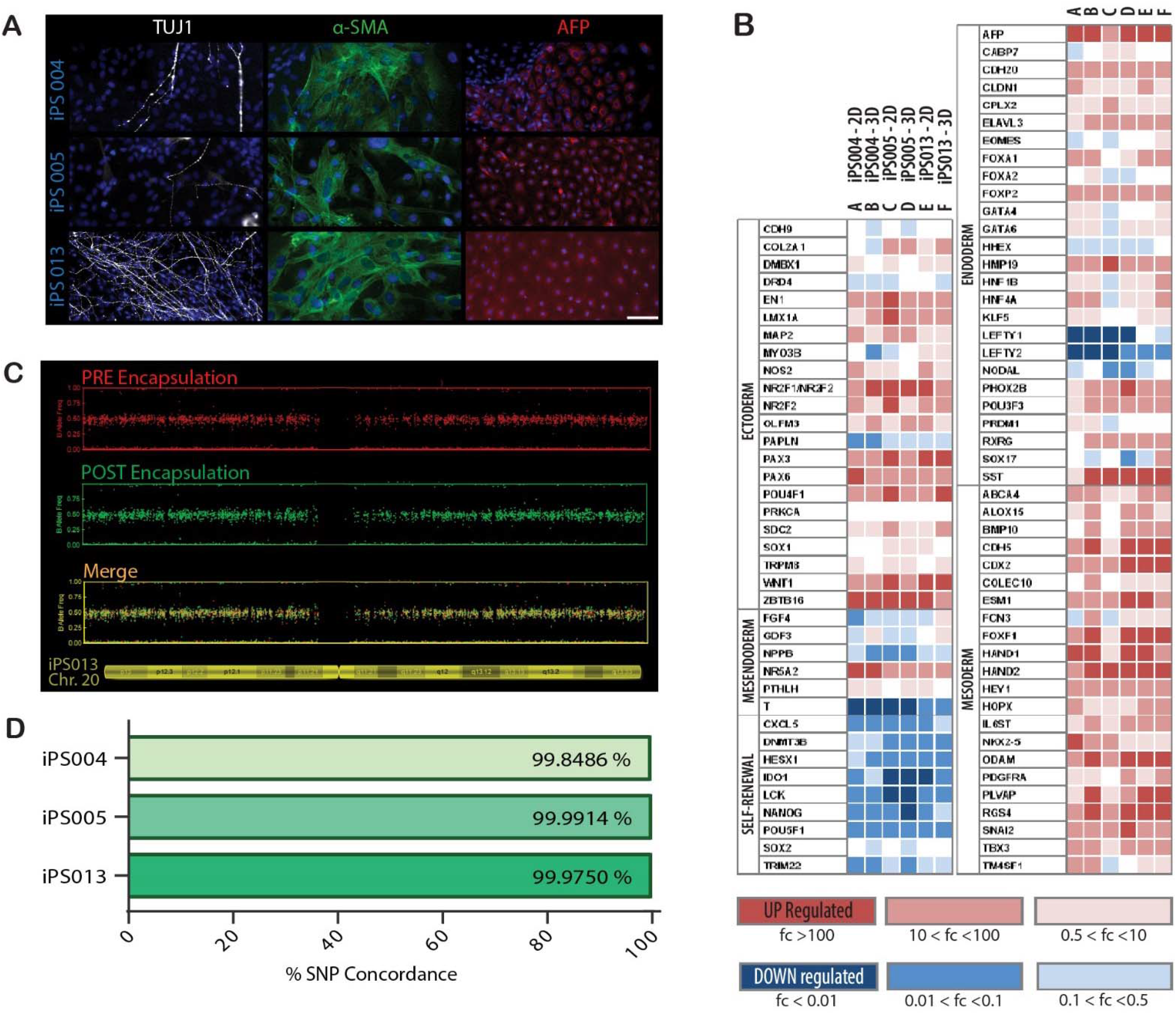
Maintenance of pluripotency and genomic integrity in 3D hPSC colonies. (**A**) Microscopy images of immunohistochemistry-based trilineage assay of 3 iPSC lines with 4 stainings: TUJ1 (White, early ectoderm), α-SMA (green, early mesoderm), AFP (Red, early endoderm), Dapi (Blue). (**B**) Scorecard^TM^ differentiation assay comparing 3 iPS cell lines expanded in 2D and 3D encapsulated hPSC colonies. (**C**) Comparison of high-resolution SNP arrays before and after one-week of encapsulation for iPS013 cell line : Zoom on chromosome 20 for pre-encapsulation (red) and post-encapsulation (green). The merge (yellow) is shown to highlight the absence of duplications and deletions. (**D**) Quantitative analysis yielding genotype SNP concordance before and after one-week of amplification as encapsulated 3D colonies for 3 cell lines.

To further quantify the differentiation potential after 3D culture within capsules, we used qPCR Scorecard^TM^ assay to evaluate the transcription profile of the cells obtained in the trilineage assay (Fig. 3B). The set of 94 previously validated qPCR markers of self-renewal, ectoderm, mesendoderm, mesoderm and endoderm (*35, 36*) was used to compare standard 2D culture and 3D culture-in-capsules. Fig. 3B shows that, for a given marker and a given cell line, there is a striking similarity between the transcription signatures in 2D and 3D culture conditions, indicating that pluripotency assessed as the *in vitro* differentiation capability of hiPSC clonies is definitely not altered in our encapsulated 3D colonies. A pooled analysis by germ layer (Fig. S4) confirms similar differentiation profile between 2D and 3D stem cells.

Finally, to control genomic integrity of the 3D colonies, we performed high resolution SNP (single nucleotide polymorphism) arrays before and after amplification within the capsules (Fig. 3C and Fig. S5) (*37*). Comparative SNP analysis showed the absence of aneuploidies, deletions or duplications, as evidenced by the superimposable karyotypes. The high degree of SNP concordance (>99.8% for all hiPSC lines) before and after encapsulation confirms cell line identity (Fig. 3D).

### Comparative growth of 3D hiPSC colonies at the scale of a single capsule, in a static suspension and in bioreactors

To evaluate whether the strategy to produce 3D colonies in ECM-coated capsules impacts the growth and expansion rates of hiPSC, we performed a series of sytematic experiments to probe the cell growth kinetics. The standard 2D cell cultures were taken as a control. Since the amplification factor is defined as AF=N(t_0_+Dt)/N(t_0_), where N(t_0_) and N(t_0_+Dt) are the cell numbers at the initial time t_0_ and t_0_+Dt respectively, direct cell counting at day 6 after passaging give a mean AF_2D_(t=6 days)∼13. Since, by definition, AF(Dt)=2^Dt/PDT^, with PDT the cell population doubling time, by pooling amplification factors at different harvesting time, we obtain a mean PDT_2D_=34h ±5 hours for iPS C line, which falls within the range of data reported in the literature (*14, 15, 38*). Then, in order to characterize the growth of individual encapsulated 3D hiPSC colonies, we cultured them in 35 mm petri dishes (typically as few as 10 capsules in a volume of medium ∼5ml, permitting to conserve the same medium for the whole course of the experiment without any risk of nutrient depletion and acidification). We performed time-lapse phase contrast imaging over a one-week period. We assume that cell volume remains constant, which allows us to derive *AF_capsule_(Dt)= V(t_0_+Dt)/V(t_0_)* by measuring the volume of the cyst *V(t)* from image analysis: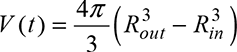 where *R_in_* and *R_out_* are the average internal and external radii of the cyst (see notations on Fig. 4A). Figure 4B shows the evolution of AF_capsule_ as a function of time for individual encapsulated 3D colonies. One immediately observes that *AF_capsule_(t=7 days)*= 212. Additionally, since AF increases exponentially as 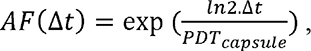 one finds *PDT_capsule_* = 22±1 hours.

**Fig. 4.**
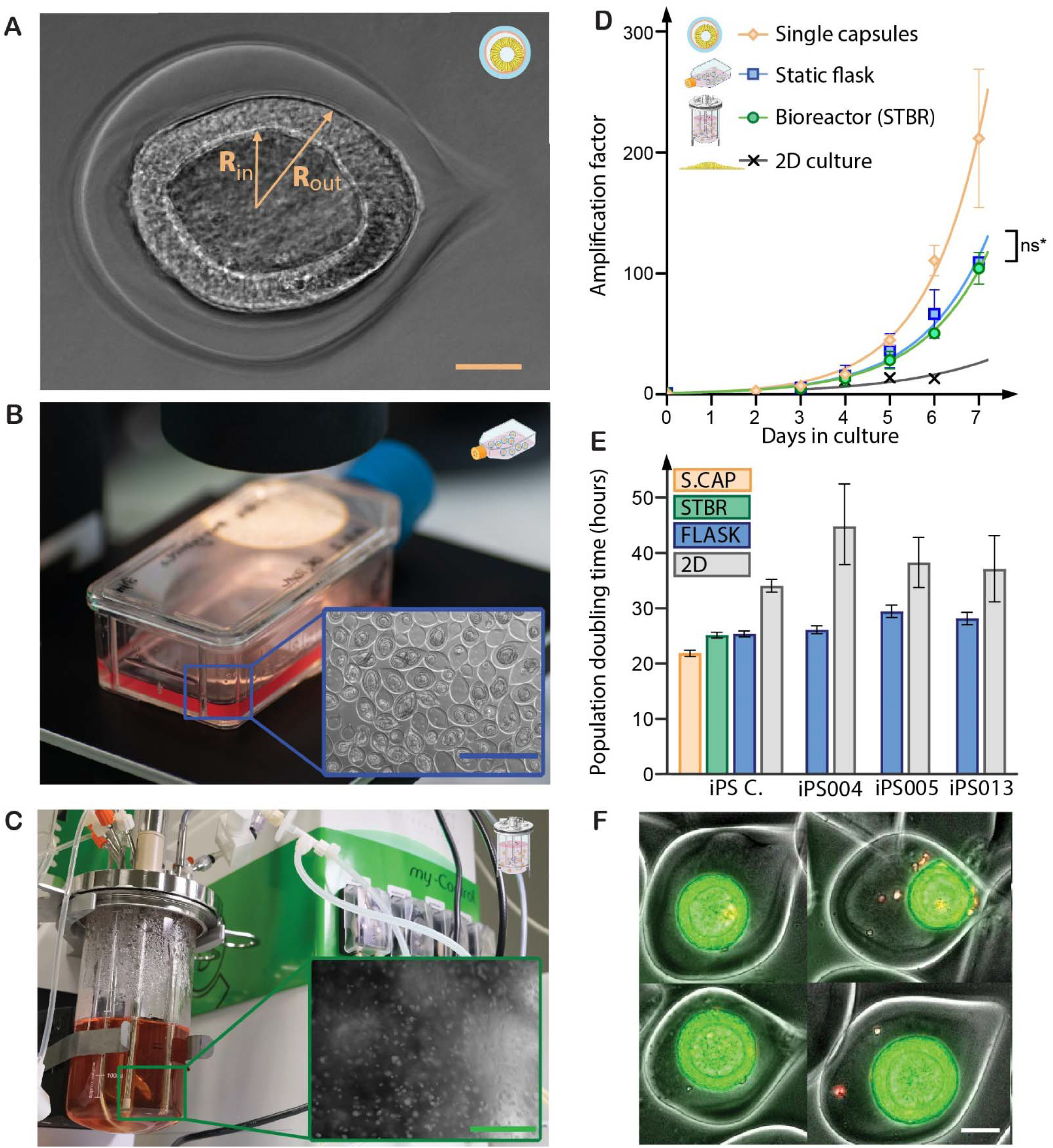
Amplification of 3D hiPSC colonies at the scale of a single capsule, in a static suspension and in bioreactors. (**A**) Micrograph of a 3D hPSC colony in a capsule and notations used in the next for the internal and external radii of the cyst. (**B**) Static suspension culture of encapsulated 3D hPSCs. Insert : Phase contrast image showing 3D hiPSCs colonies in capsules. Scale bar is 1000 µm. (**C**) Stirred suspension culture of encapsulated 3D hPSCs in a 500 ml benchtop STBR. Insert : Picture of the flowing capsules in the bioreactor. Scale bar is 4 mm. (**D**) Amplification factor as a function of time for single capsules (orange, n= 6), static culture (blue, n=2), stirred culture in a benchtop (volume 500ml) bioreactor (green, n=2), and conventional 2D cultures (grey). Last points in the graph correspond to the harvest time. (**E**) Population doubling time of encapsulated hPSC colonies in single capsules (S-CAP, orange), in a flask (FLASK, blue, n=42) and a benchtop bioreactor (STBR, green, n=2) and in standard 2D cultures (2D, grey). Error bars represent the standard error of the mean. (**F**) Fluorescence microscopy image of 4 representative encapsulated 3D hPSC colonies stained with Live/dead (green/red)). Scale bar is 100 µm. All data shown here were obtained with iPS C line, except panel 4D that collects data for the 4 available cell lines.

With the perspective of scaling up the production of hiPSCs, we also investigated how the growth of encapsulated 3D hiPSC colonies was impacted when cultured in conditions of i) static bulk suspension in standard T-Flasks (Fig. 4B) and ii) stirred suspension in a benchtop bioreactor (Fig. 4C). STBRs are the most common bioreactors used to culture biological agents for biotechnological applications. Besides ther capacity to monitor and adjust the pH and oxygen partial pressure and to refresh the medium, the mechanical agitation provided by the impellers allows better fluid mixing and oxygen transfer ability as compared to static suspension (*39*). However, the drawback may also be that the shear stess induced by the impellers was shown to cause deleterious effects such as cell death or decrease in cell growth in aggregate- or microcarrier-based cultures (*40, 41*). Practically, we loaded capsules from a same batch in T-flasks and in a STBR at the same initial density. The bioreactor impeller rotational speed was set to 150 rpm, which is sufficient to maintain medium homogeneity and avoid capsule sedimentation. We could not detect any change in the shape of the capsules and 3D colonies under these stirring conditions. Then, after dissolution of the alginate shell and dissociation of the cysts, we counted the stem cells in both static and stirred culture conditions to derive the amplification factors AF in time. We found *AF_flask_(7 days)*= 109 ± 6 and *AF_bioreactor_(7 days)*= 104 ± 18 (Fig. 4D). From the values of the characteristic times for the exponential variation of AF, we could calculate, as explained above, *PDT_flas_*_k_= 25± 6h and *PDT_STBR_*=25±3 h (Fig. 4E). Three remarks can be made. First, these PDT values are significantly lower than the ones derived from 2D cultures, indicating again than the expansion is greatly improved in 3D, as evidenced by the low number of dead cells in capsules (Fig. 4F) as compared with 2D colonies (Fig. S6). Second, the AF values are about twice as low as the one derived from the masurements at the single capsule level. Third, the absence of statistical difference between the two culture systems suggests that, while the impeler-induced shear stress does not affect cell viability, stirred suspension culture in a benchtop bioreactor with expected better homogenization does not enhance the expansion under the experimental conditions selected here.

Additionally, we performed flow cytometry analysis and found that more than 92% of the cells are positive for SOX2, NANOG and OCT4. Stemness thus remains high and similar between static and stirred cultures (Fig. S7), indicating that, by contrast with previous reports (*42*), shear stress does not trigger the differentiation of hiPSC colonies grown in hollow capsules. Finally, in order to confirm that the amplification factors reported above are not hiPSC line specific, we carried out the same series of experiments for the other 3 cell lines in static culture (Fig. 4E). Not only are the differences between cell lines not significant, but their PDT in 3D is also found very close to the value derived for the commercial hiPSC line that we have extensively investigated in this section, i.e. *PDT_flask_*= 27± 2 h by averaging over all 3 cell lines.

### Optimized culture conditions: Impact of capsule size and oxygen tension

As shown above, the static or stirred batch cultures of hiPSC colonies in capsules yielded amplification at day 7 about twice as low as the one measured at the single capsule level. Although this difference only corresponds to a 3h difference in PDT, we sought to address this issue and find out solutions to further improve the amplification of hiPSC in batch for large-scale production. We investigated the impact of two possible parameters.

First, we tested whether hiPSC amplification depends on the cell seeding density. The most obvious way would be to increase the volume fraction of cells in the core solution loaded to the microfluidic injector. However, this would lead to earlier and more frequent harvesting. Instead, we pursued a different strategy. We kept the cell density constant but increased the size of the capsules by changing the size of the nozzle (*28*). Doing so, for a given volume of the encapsulation cell suspension, the number of produced capsules is indeed reduced by *(R_big_/R_small_)^3^* but the mean number of cells per capsule, λ, is increased by the same fold. In the context of sparse distribution, Poisson statistics applies and the generation of 300 µm in radius capsules instead of 200 µm leads to an increase in λ by about 3 fold. The immediate consequence is that the probability to have capsules containing no cell is decreased from about 8% to negligible (∼0.03%). But, more importantly, the probability to have capsules with only one cell that may die or exhibit some lag phase before proliferation goes from 20% to 0.3%. High occurrence of capsules loaded with only one cell is expected to lower the effective amplification factor and thus to increase the doubling time of the cell population. Fig. 5A shows representative phase contrast images of 300 µm in radius capsules filled with hPSC colonies. Noteworthily, most capsules contain several cysts, suggesting that cyst growth was nucleated from several cell aggregates. More quantitatively, after monitoring the growth kinetics in benchtop STBR, derivation of the time constant reveals shorter doubling time in big capsules: *PDT_big_* =22h±1h < *PDT_small_*=25h±1h (Fig. 5B).

**Fig. 5.**
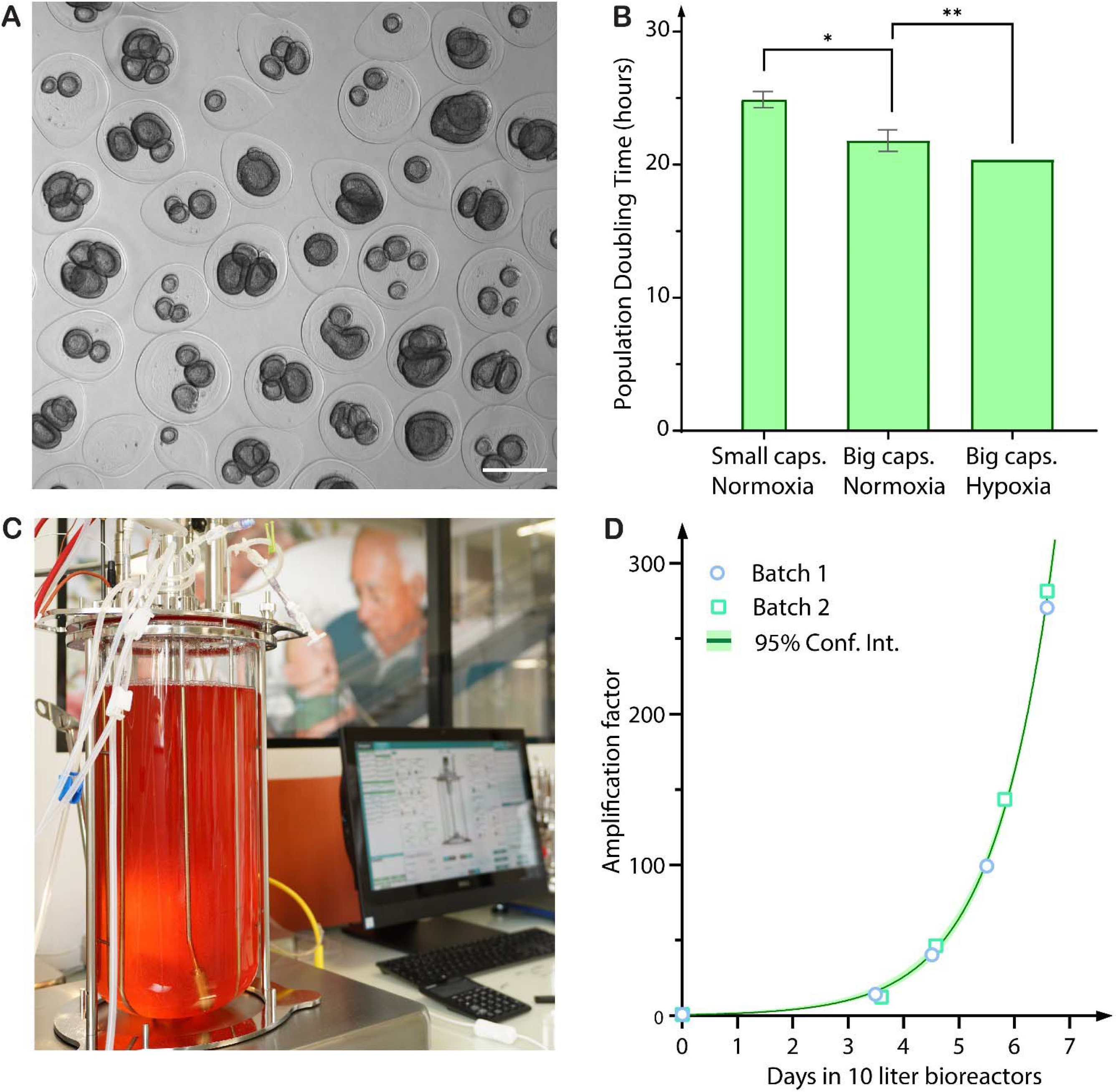
Impact of capsule size and oxygen tension on hPSC amplification and scalability in stirred tank bioreactors. (**A**) Phase contrast image showing 3D hiPSCs colonies in capsules referred to as “big” in the main text (with an average radius of 300µm). Scale bar = 500µm. (**B**) Population doubling time of encapsulated 3D hiPSCs colonies cultivated in benchtop bioreactors by varying the size of the capsules and the oxygen tension conditions (normoxic versus hypoxic). Mean and standard deviation (* p<0,001 and ** p <0,01). (**C**) Picture of a 10 liter industrial stirred tank bioreactor used to test the scalability of the stem cell capsule culture system. (**D**) Graph of amplification factor of hiPSCs grown in 10 liter bioreactors over a week, in ‘Big capsules’ and hypoxic conditions; Data were obtained from 2 independent batchs and from 2 independent encapsulations. Light green band shows the 95% confidence interval of the fitting curve.

Second, we pursued along our physiomimetic approach. Among all factors that make a stem cell niche, we have already recapitulated interactions with the ECM. However, untill now, we have omitted to consider oxygen tension, which is known to be naturally low in developping embryos (*43*). This low level of oxygen was further shown to be key to reduce mutation rates and epigenetic alterations (*44, 45*) as well as to improve the expansion rate (*46*) while reducing the probability of unwanted differentiation (*47*). We thus performed the same culture experiments in big capsules (300 µm radius) by decreasing the dissolved oxygen level (DO) from 100% to 20%. (see Materials and Methods section). Under these hypoxic conditions, the population doubling time derived from the growth kinetics was found to be 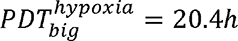 (Fig. 5B), i.e, slightly but significantly shorter than in normoxic conditions. Remarkably, this value is better than the one found at the single capsule level in normoxia, suggesting that the optimization of capsule size and oxygen tension allowed to upscale the production of hPSCs in a bench-scale bioreactor without any degradation of the expansion efficiency.

In order to assess whether this protocol is not only theoretically scalable but can be truly upscaled to an industrial level, we carried out a final experiment in duplicate in a 10 L STBR (Fig. 5C, S8 and Movie S5) by keeping all other parameters constant. Figure 5D shows the the expansion-fold as a function of time. The two curves from these two independent experiments are superimposable and we found *AF_10L STBR_(6.5 days)*=277, corresponding to a doubling time *PDT_10L STBR_*=19.6 h.

These data were obtained without passaging. In order to assess the robustness of the approach for a seed train of passaging and demonstrate that the technology can be integrated into a classical cell therapy production, we performed serial passaging. At harvest after about 7 days, the capsules were dissolved, the 3D colonies were dissociated and cells re-encapsulated following the same protocol (Fig. 6). We found that expansion-fold and stemness were preserved over two consecutive encapsulations within 14 days (Fig. S9).

**Fig. 6.**
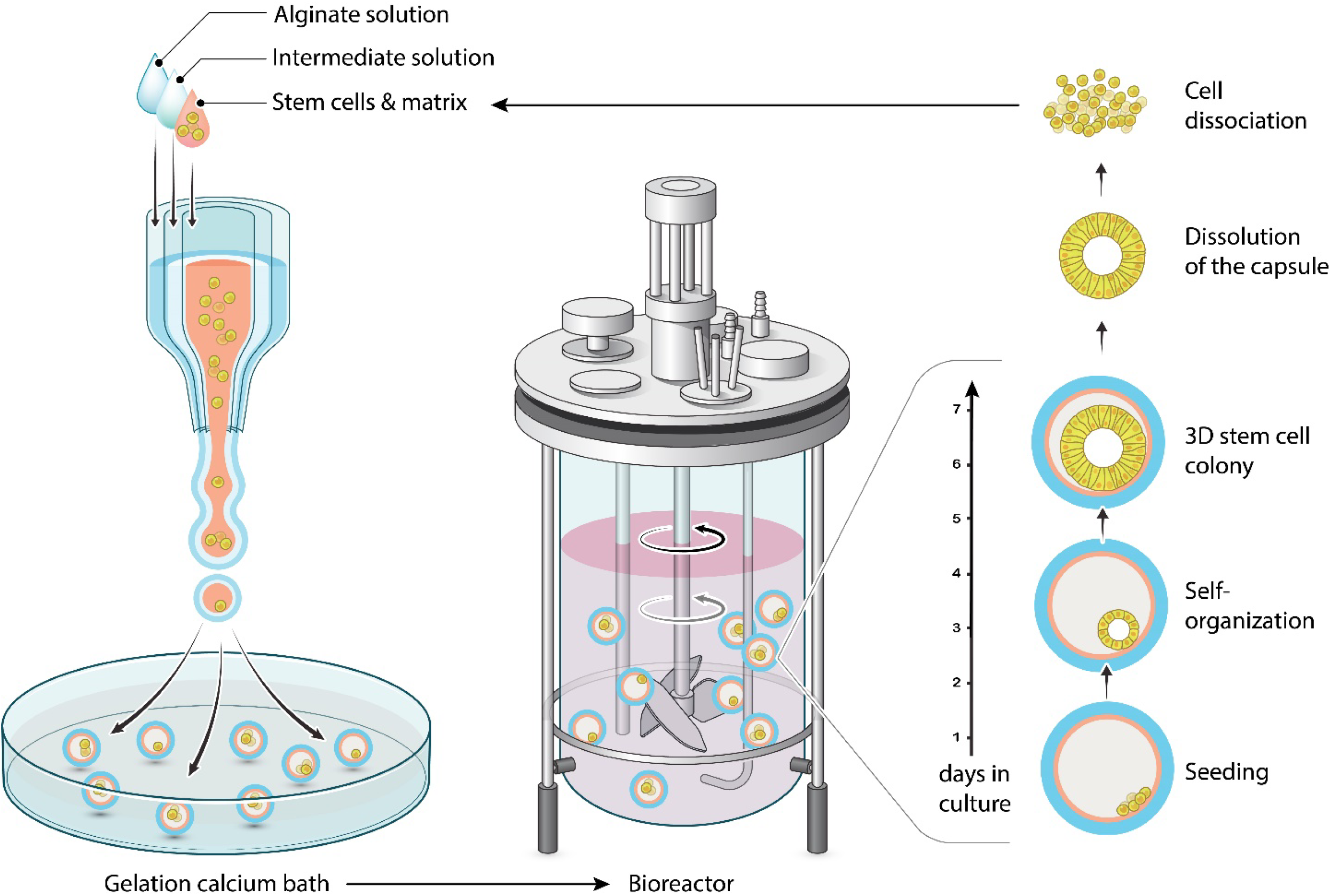
C-STEM pipeline: encapsulation and scale-independent culture of encapsulated 3D hPSC colonies in bioreactors. After encapsulation using the microfluidic extrusion technique (left panel), hPSC in matrix-laden capsules are transferred to suspension culture in a bioreactor (middle panel). Under controlled conditions provided by the bioreactor, hPSC cells self-organize into cysts which are protected by the capsules. These growing 3D colonies are harvested and dissociated after capsule dissolution. Subsequent cell suspension may then serve for another encapsulation and expansion.

## DISCUSSION

In this work, we have developed an *in vitro* culture system for hiPSCs, that we named C-STEM (Fig. 6) and combines the benefits of biomimetic 3D culture and scalable bioreactor-based production. By contrast with other approaches using scaffold embedding in bulk matrix (*48*) or hydrogel beads (*49, 50*), our hollow capsules allow in-situ engineering of stem cell niche-like microenvironment. With biological and topological cues driving 3D self-organization of hiPSCs colonies that are reminiscent of epiblasts (*51*). While cell-cell interactions are not impaired due to the absence of enwrapping scaffold, the presence of the shell also provides mechanical protection against impeller damage and turbulence-induced so-called Kolmogorov eddies during stirring (*52*). Besides exploiting both the biomimetic and protective properties of the developed C-STEM platform, we have finally refined the culture conditions by optimizing the initial mean number of cells per capsule and the oxygen tension. Combination of all these critical factors allowed us to upscale the production of hPSCs and demonstrate that the amplification efficiency is scale-independent. In particular, we could reach 282-fold amplification in 6.5 days in a 10 L stirred-tank bioreactor.

To the best of our knowledge, this level of both amplification and scalability is unmatched in the field (*15, 53*). This technical and quantitative tour de force is actually related to the higher cell viability obtained in our hollow capsules as compared with 2D cultures or other 3D suspension cultures. Since the amplification factor *AF(t)* of a culture system is given by *AF(t)* = 2*^t/PDT^*, where PDT is the population doubling time and accounts both for cell division and cell death, we may rewrite it as *AF(t) = 2^(k_+_-k_-_)t^* where k**_+_** and k**_-_** are respectively the division and death rate of cells. Thus, the upper theoretical limit for AF is obtained for k**_-_**=0 (i.e. infinite death time), which then yields a minimal *PDT_min_* value, equal to k**_+_**. Quite surprisingly, the measurement of k**_+_** or the duration of the cell cycle t**_+_**=1/k**_+_** of hiPSC has been overlooked in the literature. The sole report we are aware of arises from a recent work in which cell cycle kinetics was tracked using a rainbow reporter in primed pluripotent stem cells (*54*). The obtained cell cycle duration was found to be t**_+_**∼14 h. While the question of “gemellarity” between embryonic stem cells (ESC) and hiPSC is still under debate (*55*), this value is also consistent with previous estimates of t**_+_**=11–16-h for the cell cycle duration in human and mammalian ESC (*56–58*). By assuming that the duration of the cell cycle in the 3D cyst topology is identical to the value obtained in 2D cultures with AF(t=6.5 days)=282 corresponding to PDT=19h, we find that the doubling time of the encapsulated hiPSC colonies is only 5h±2h longer than the intrinsic cell cycle duration. The difference corresponds to a death rate k_-_=1/53h^-1^. By comparison, in the seminal Yamanaka’s paper (*14*), doubling times in 2D hiPSC colonies of about 45 h indicate that k**_-_**=1/20h^-1^. More meaningful than the death rate k**_-_**, the fraction of dead cells can be roughly estimated as 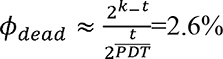 at t=6.5 days, while cell counting gives 1.30% in bioreactors and 1.97% in flasks (Fig.S6). In 2D cultures, we measured a fraction of dead cells of the order of 12% at harvest, even though this value under-estimates the cumulated mortality which is drastically impacted by cell passaging and confluency (*59*). Further expansion improvement is theoretically within reach by vanishing the cell death rate. Taking again t**_+_**∼14 h for the cell cycle length of hiPSC, the glass ceiling is calculated to be *AF_max_(t=6.5 d)*∼2200. However, the value of t**_+_**∼14 h cannot be taken as granted. For instance, a cell density-dependence on the proliferation rate has been reported.

Indeed, hPSC exhibit a decelerated proliferation due to a prolonged G1 phase as cell density increases (*60*). Similarly, smaller expansion folds were observed when the inoculation density either as single cells or pre-clusters in a synthetic hydrogel exceeds 10^6^ cells/mL (*48*). We may then anticipate that the actual average cell cycle duration could be longer than the one reported above upon single-cell lineage tracking in small colonies of hPSCs. As a consequence, even though future work should aim at a rigorous *in situ* measurement of the division rate within the encapsulated hiPSC cysts, we cannot exclude that the unprecedented hiPSC expansion rates reported in this work are approaching the glass ceiling.

Among other specificities of the C-STEM technology, we mentioned the protective role of the alginate shell and the scale-independence of the culture conditions. By contrast with other suspension cultures that need to design specific low-shear impellers (e.g. the vertical-wheel bioreactor, (*61, 62*), or to add shear-dampener polymers (e.g. pluronics, (*24*)) in order to avoid stirring-induced cell damages, our capsules permit the use of standard industrial scale bioreactors.

However, all the benefits cannot be assigned to this sole shielding effect. Indeed, previous works had already proposed to embed either hESC aggregates or microcarriers within hydrogel beads (which are referred to as capsules in these original works) (*50*) to improve cell viability. Nonetheless, expansion rates were not reported to be larger than 10 in 19 days. Similarly, two recent works describe stem cells encapsulation in hollow capsules (*63, 64*). However, the absence of ECM leads to the formation of aggregates and a modest amplification (estimated to 70-fold in 8 days from the size of the encapsulated spheroids). We thus propose that the significant amplification increase obtained with the C-STEM technology mostly originates from the stem cell niche-like environment that is engineered within each capsule and that drives hiPSC multicellular organization into cysts.

Interestingly, numerous studies have recently proposed biomimetic controllable environments that can be used to develop hPSC-based embryo models and more specifically epiblast models (*22, 65, 66*). In all cases, these 3D stem cell niche mimics drive PSCs self-organization, luminogenesis, and polarization into pseudo-stratified epithelia (*19–21*). This cyst configuration, which seems to be key in developmental processes, may be regarded as an optimized configuration for hPSC expansion with minimal loss of viability (*67*). First, it is well accepted that 2D hiPSC cultures exhibit intra-colony heterogeneities in pluripotency marker genes (*68*), viability (*69*) and cell morphology (*70*), which are very striking between the center and the edges of the colonies. In this respect, the closed spherical symmetry of a cyst intrinsically suppresses the “center-edge phenotype” and may result in more homogeneous cell population (*71*). Additionally, whereas cellular crowding or compaction are known to inhibit proliferation or even trigger apoptosis via caspase-dependent mechanisms (*72*) a cyst configuration is less prone to stress building in bulk due to the presence of a lumen. Similarly, fast proliferation rate may contribute to stress relaxation and reduce cell extrusion occurrence as observed in epithelia under compression (*73, 74*). Besides mechanical stress, chemical stresses are known to increase the mutation rate (*44, 75*). The use of bioreactors with precise adjustment of physioxia, pH, lactate, glucose and nutriments supply is thus instrumental and could be optimized beyond the present achievement.

Finally, relying on the observation that chromosome segregation fidelity is unambiguously higher in native contexts of epithelia of primary cells (*76, 77*), it also becomes tempting to speculate that, beyond the gain in amplification it provides, the preserved histology of our *in-vitro* epiblast-like colonies could also be beneficial to the maintenance of the genetic integrity (*78*).

In summary, our work has shown that hollow alginate capsules with reconstituted niche-like microenvironement can promote the formation and growth of 3D hPSC colonies and provide the necessary protection for scaling up the production in stirred tank bioreactors. Self-organized encapsulated epiblast-like colonies seem to be instrumental for optimal expansion by preserving stem cells physiological properties. We have demonstrated that our biomimetic stem cell platform C-STEM can deliver unprecedented scalability and we anticipate that cell quality is preserved on the basis of extremely high viability, which is taken as a primary signature of cell fitness. Future works should focus on assessing the hPSCs quality in *in vivo*-like culture systems, since the emergence of mutations during culture may be the last limitation to overcome for cell therapy bioproduction.

## MATERIALS AND METHODS

### Ethics statements

The generation, use and storage of hiPSCs were performed with approval from the “Comité de Protection des Personnes” (CPP) Ile de France (DC 2015-2595 and 2016-A00773-48).

### Human pluripotent stem cell lines

Throughout the present work, we used 4 hiPSC lines. Among these, 3 hiPS cells, namely IMAGINi004-A (referred to as iPS004), IMAGINi005-A (iPS005) and IMAGINi013-A (iPS013) were derived from peripheral blood mononuclear cells (PBMC) according to the protocol described in (*34*). Briefly, PBMCs were transduced using the CytoTune-iPS 2.0 Sendai Reprogramming Kit (ThermoFisher Scientific) following the manufacturer’s instructions. After 2-3 weeks, colonies were manually picked and expanded at least 10 passages. The 4^th^ hiPSC line is a commercial line from ThermoFisher: Gibco^TM^ episomal hiPSC line (A18945) generated using cord blood derived CD34+ progenitors with 7 episomally expressed factors (Oct4, Sox2, Klf4, Myc, Nanog, Lin28, and SV40 T). This commercial cell line is referred to as iPS C. For the sake of availability, in order to allow other groups to reproduce our findings, all experiments reported here were performed with iPS C, except for those that are described in Figures 2F, 3A,B,D 4E and Figures S4 S5, which were carried out to demonstrate that the findings were not cell line-dependent.

### 2D hiPSC culture

All hiPSC lines were maintained on Matrigel (Corning Ref. 354234) and cultured in mTeSR1 medium (StemCell Technologies 85875). Cultures were fed daily, passaged with an enzyme-free reagent, ReleSR (StemCell Technologies 05873) every 3-6 days (around 80% confluency) and replated as small clusters (between 100 and 200 µm) at a density of about 5000-10000 cells/cm^2^. Cells were cultured at 37°C in a humidified atmosphere containing 5% CO2.

### 3D hiPSC Encapsulation

Prior to encapsulation, 2D stem cell colonies were detached using ReLeSR for 1 minute and dissociated into a near single cell solution using Accutase (Stem Cell Thechnologies 07920). HiPSCs were then mixed in a 50/50 volume ratio with Matrigel at 4°C to keep the suspension in a liquid state. The final concentration of cells in the cell/matrix solution was thus between 0.4-1.0×10^6^ viable cells/mL, referred to as the encapsulation density. The encapsulation system is similar to the one described in (*29*). In brief, ethylene tetrafluoroethylene (ETFE,) tubings are connected to the three inlets of a 3D printed (using the DLP Micro Plus Hi-Res printer from EnvisionTEC) microfluidic co-laminar flow device. An extruded and polished glass microcapillary tip (of diameter ∼100 µm for most experiments reported in this work, at the exception of those shown in Fig. 5A-B that were carried out with a nozzle diameter of 150 µm) is glued to the outlet of the nozzle for a better control of the flow. The cell/matrix suspension is loaded into the inner channel of the 3-way device, which is kept refrigerated thanks to an in-line cooling system in order to prevent premature gelation of Matrigel. A solution of sodium alginate (Novamatrix Pronova SLG100, 0,25 g #4202101 at 2% in distilled water) is injected into the outer channel. To prevent alginate gelation within the microfluidic device due to calcium release from the suspended cells, a calcium-free solution (Sorbitol 300mM, Sigma-Aldrich 85529) is used in the intermediate channel of the co-extrusion chip and serves as a barrier against calcium diffusion. Typical flow rates for the 3 solutions were on the order of 120 ml/h for all three channels: (alginate solution, the sorbitol solution and the cell+matrix suspension). At these rates, the composite solution forms a liquid jet that fragments into droplets (of about twice the size of the nozzle) due to the spontaneous Rayleigh-Plateau instability. To avoid subsequent coalescence of the train of droplet, an alginate charging part and a copper ring are connected to a high voltage (2000V) generator are introduced. A high-speed camera (PHANTOM VEO 1310L) was used to visualize droplet formation and splay (Fig. S1 and Movie S1). When the composite droplets contact the collecting calcium bath (at 100mM), the outer layer of alginate readily gelates. As a consequence, the inner cell/matrix solution remains entrapped inside a closed, spherical and permeable micro-compartment. Within 1 min following encapsulation, capsules are rinsed with medium (DMEM) to reduce the basal calcium concentration. Finally, ther are transferred to suspension culture medium.

Re-encapsulation was performed by dissolution of alginate shells using short rReleSR rinse, followed by cell dissociation using TrypLE (Trypsin-based, dissociation enzyme, ThermoFischer) for 20 minutes at 37°. Then the obtained cells were processed following the classic encapsulation protocol.

### 3D stem cell suspension culture in static T-Flasks or bioreactors

Static suspension cultures of encapsulated hiPSC were carried out using T-Flasks (from 5 to 30 ml) maintained in a cell culture incubator at 37°C and 5% CO_2_. The medium (mTeSR1) was supplemented with 10µM Y-27632 for ROCK inhibition only during the first 24 hours of culture. From culture day 3, the medium was exchanged every day as described hereafter. The contents of the T-Flasks were transferred into Falcon Tubes. After capsules sedimentation (within a few minutes), the supernatant was removed and replenished as the capsules were transferred back into a T-Flasks. The volume of culture medium was kept constant for the first 4 days of culture (∼ 4× the capsules volume). Then, the volume was steadily increased every day in order to maintain a cell concentration bellow 10^6^ cells/mL.

Stirred suspension cultures were performed in different bioreactors. For all experiments reported in Fig. 4 and Fig. 5A-B, we used benchtop STBRs, including a 30 mL (Minibio, ABLE® Bioreactor Systems) or 500 mL bioreactors (Applikon Biotechnology & Global Process Concept). For the experiments reported in Fig. 5C-D, we used a 10 liter-scale bioreactor (Global Process Concept).

In all cases, the bioreactors were inoculated with 15% capsule-to-medium volume. The bioreactor culture starts at a volume representing 30% of the final working volume. At Day1, the medium was replaced with fresh medium without ROCK inhibitor. From day 2 to 5, the culture is performed in a fed-batch mode up to the final working volume (*39*). Then, we switched to repeated-batch mode, where 90% of the media was daily renewed to maintain sufficient nutriment supply. The final capsule-to-medium volume was 4.2 ± 0.3% and the pH was maintained at 7.2 ±0.2.

Dissolved Oxygen (DO) level is calibrated at 100% in starting conditions by injecting air into the bioreactor headspace. During the run, the oxygen level is monitored and controlled. In normoxic conditions the oxygen is controlled at 100% while in hypoxic conditions the set point is at 20% DO. Oxygen level is regulated by sparging nitrogen and/or air to maintain the set point. 10L scale bioreactors were regulated in hypoxic conditions. During one week of culture, the stirring speed is set at 150 rpm that is sufficient to keep capsules resuspension and bioreactor homogeneity along the run.

### Time-lapse microscopy of encapsulated cyst growth and image analysis

Time-lapse microscopy was performed using a Nikon Biostation IM microscope with a 10x objective. Capsules containing hiPSCs were transferred to a 35 mm Petri dish 24 hours after encapsulation. Approximately 10 to 20 capsules were placed in the petri dish containing 5 mL of fresh Y-27632-free mTeSR1 medium. Cyst growth was monitored over 7 days. Practically, images were taken every 6 to 10 minutes at preselected Z-focal planes to ensure acquisition at proper focus in case of undesired drift Image analysis was performed using ImageJ. The external and internal effective radii of the cysts, R_out_ and R_in_, were measured from the equatorial corresponding cross sections S according to R_out,in_=(S_out,in_/π)^1/2^ after applying appropriate bandpass filters and thresholds. The volume V of the cells was calculated as V=4π/3(R_out_ ^3^-R_in_ ^3^). Capsule circularity was defined as C=a^2^/b^2^, where a and b are the short and long axes of the approximated ellipse.

### In vitro trilineage differentiation

Small cell clusters were collected from hiPSC cultures (2D or decapsulated-dissociated 3D hiPSCs colonies) and transferred into low attachment dishes (Corning, Ultra-low attachment 6 well plate). Three-dimensional aggregates of cells that are an amalgam of the three developmental germ layers (Embryoid bodies) are obtained and cultured in suspension for 7 to 9 days with DMEM/F-12 medium containing 20% pluriQ Serum Replacement (GlobalStem), 1% non-essential amino acids, 1% penicillin-streptomycin and 0.2% β (ThermoFisher Scientific) in a humidified atmosphere containing 5% CO2. Culture medium was refreshed every two days. EBs were then collected for RNA analyses or transferred onto gelatin-coated dishes for 1 week. For immunocytochemistry analysis, cells were fixed with 4% paraformaldehyde for 15 min at room temperature (RT). After washing in PBS/1%BSA blocking solution for 1 hour, cells were incubated overnight at 4°C with primary antibodies, washed 3 times in PBS, and incubated with secondary antibodies for 2 hours at RT. Antibodies were diluted in PBS/1%BSA/0.1%triton solution. The list of antibodies used in this work and their origin are listed in Table S1. Nuclei were stained with a DAPI solution. Immuno-fluorescence staining was analyzed using the Celena S™ Digital Imaging System (Logos Biosystems).

### RNA extraction and RT-PCR analyses

Total RNA was extracted using the RNeasy Mini Kit (Qiagen). cDNA was synthesized using a high capacity cDNA RT kit (ThermoFisher Scientific) from 1μg of total RNA. The expression of pluripotency markers as well as the trilineage differentiation potential of the cells were evaluated by TaqMan® hiPSC Scorecard™ assay according to the manufacturer’s protocol. This scorecard compares the gene expression pattern of key pluripotency and germ lineage markers relative to a reference standard that consists of 9 different human ES and iPS lines. Data analysis was performed using the cloud based TaqMan® hiPSC Scorecard^TM^ analysis software.

### DNA isolation, genomic stability and authenticity analysis

DNA isolation was performed using the PureLink™ Genomic DNA Mini Kit (Invitrogen). Molecular karyotype was performed using an Infinium Core-24 v1.2 Kit (Illumina) containing 300000 SNPs. Data were analyzed with BeadStudio/GenomeStudio software (Illumina). The percentage of SNP concordance between iPSC samples before and 7 days after encapsulation was assessed for the 3 derived iPS cell lines. SNP files of all samples were extracted from genome studio software. The percentage of concordance between two paired samples (before and after encapsulation) was evaluated by comparing the genotype of each informative SNP (Fig. S5)

### Flow cytometry analysis

The hiPSCs colonies were dissociated with Accutase for 10 minutes at 37°C for 2D cultures or with TrypLE Select (ThermoFisher Scientific 11598846) for 30 minutes at 37°C for 3D cultures after capsule removal. Then, cells were fixed and permeabilized using the Transcription Factor Staining Buffer Set (ThermoFisher 11500597). Cells were suspended in the permeabilization buffer at a density of 0.5-1×10^6^ cells in 100 μl and incubated with the specific antibodies or isotype controls (Table S1) for 45 minutes at room temperature in the dark. Cells were washed twice with the staining buffer and analyzed using either BD Canto II (at the TBMCore CNRS UMS 3427 – INSERM US 005) or a BD Accuri^TM^ C6 plus and data was post-processed with FlowJo software.

### Cell growth and viability analysis

Cell counting was performed using the Nucleo counter NC3000 or NC200 (Chemometec). Live/dead analysis was performed using CalceinAM/Ethidium homodimer-1 (ThermoFisher L3224) according to the manufacturer recommendations, and samples were imaged using either the EVOS FL or EVOS M5000 auto Imaging system (ThermoFischer).

### Immunostaining, Microscopy, and Image Analysis

For daily brightfield imaging of 2D cultures and encapsulated hiPSC cysts, a widefield EVOS FL or EVOS M5000 automated microscope was used. Encapsulated 3D hiPSC colonies were harvested for confocal microscopy at several timepoints. The alginate capsule was removed prior to fixation by incubating the samples in PBS without divalent cations for at least 5 minutes with agitation at RT. Both 2D and 3D stem cell colonies were fixed with 4% PFA for 30-60 minutes at RT in the dark. Following fixation, the samples were washed 3x with 0.1% Tween20 in PBS. A permeabilization step was done in parallel with excess PFA quenching in a PBS solution containing 0.3% Triton X-100 and 100 mM Glycine for 30 minutes, followed by 3x washing with 0.1% Tween20 in PBS. The samples were incubated in primary and secondary antibodies (Table S1) in 1% BSA + 0.1% Tween20 in PBS overnight at 4° with gentle orbital agitation, including a 3x rinsing with 1% BSA + 0.1% Tween20 in PBS after each incubation. To maintain alginate capsules during fixation and staining, the decapsulation step was skipped and all solutions (including 4% PFA) were supplemented with calcium and magnesium. All samples were imaged on either a Leica SP5 or SP8 confocal microscope (Bordeaux Imaging Center, BIC).

### Statistical Analysis

All statistical analyses were performed using GraphPad Prism8. To test significant differences between bioreactor and flask, 2wayANOVA with sidak’s multiple comparaison test was used. To compare between 2 groups, a student T-test was used. All statistical significance is reported in terms of p-values <0.05.

## Supporting information

Fig. S1. Optical measurement of the capsule production rate.

Fig. S2. Phenotype of encapsulated 3D hPSC colonies

Fig. S3. Stemness staining of encapsulated 3D hPSC colonies

Fig. S4. Scorecard quantitative comparison of gene expression profiles from trilineage differentiation assays of 2D and 3D hPSC colonies

Fig. S5. Analysis of high-resolution SNP arrays before and after C-STEM amplification

Fig. S6. Cell viability in 2D and encapsulated 3D hPSC cultures

Fig. S7. Encapsulated epiblast-like colonies resilience to hydrodynamic damages

Fig. S8. Key parameters and results of C-STEM scale-up in 10 liter bioreactor

Fig. S9. Stemness maintenance assessment of hiPSCs through 2 consecutive encapsulations

MOVIE S1. Formation and collection of capsules in calcium bath

MOVIE S2. 3D rendering of an equatorially sectioned 3D hPSC colony

MOVIE S3. Time-lapse of an encapsulated 3D hPSC colony : between encapsulation and harvest

MOVIE S4. Time-lapse of an encapsulated 3D hPSC colony : between encapsulation and lumen collapse

MOVIE S5. Encapsulated 3D hPSCs colonies cultured in a 10 liter stirred-tank bioreactor

## Funding

This work was supported in part by funding grants from European commission H2020-EIC-SMEInst (grant agreement number: 881113 C-stemGMP), iLAB2018 Bpi France, Region Nouvelle Aquitaine and Agence Nationale pour la Recherche (ANR-17-C18-0026-02). We acknowledge the Bordeaux Imaging Center, a service unit of the CNRS-INSERM and Bordeaux University, member of the national infrastructure France BioImaging supported by the French Research Agency (ANR-10-INBS-04). We also acknowledge the TBMCore facility. (CNRS UMS 3427 – INSERM US 005). We also thank Nicolas Doulet and Marion Pilorge for helping build the collaboration between Treefrog Therapeutics and the Imagine Institute.

## Author contributions

KA NL MF designed the project and supervised experiments and analysis; PC performed the experiments, analyzed the data and wrote the article; Experiments were performed both at the Imagine Institute and at Treefrog Therapeutics (TFT). AL helped performing experiments, analysing the data and writing the article; EL FM ML designed bioreactor cultures and scale-up, EL JP HW EJ MD performed bioreactor cultures; JC EW EQ CB EP contributed to 2D cultures, encapsulations, trilineage assay and SNP analysis; BG contributed to data analysis and writing.; PVL proposed a mathematical formulation for PDT and viability. PN helped analyze the growth of 3D colonies and write the manuscript. KA CR JH conceived and produced the microfluidic chips and performed high speed camera recordings.

## Competing interests

MF and KA are the founders of TFT; MF, KA, PC and PN are shareholders of Treefrog therapeutics. MF KA and PN have a patent pertaining to discoveries presented in this manuscript. Patent no: WO2018096277A1.

## Data and materials availability

All data are available in the main text or the supplementary materials.

## SUPPLEMENTARY MATERIAL

**Fig. S1.**
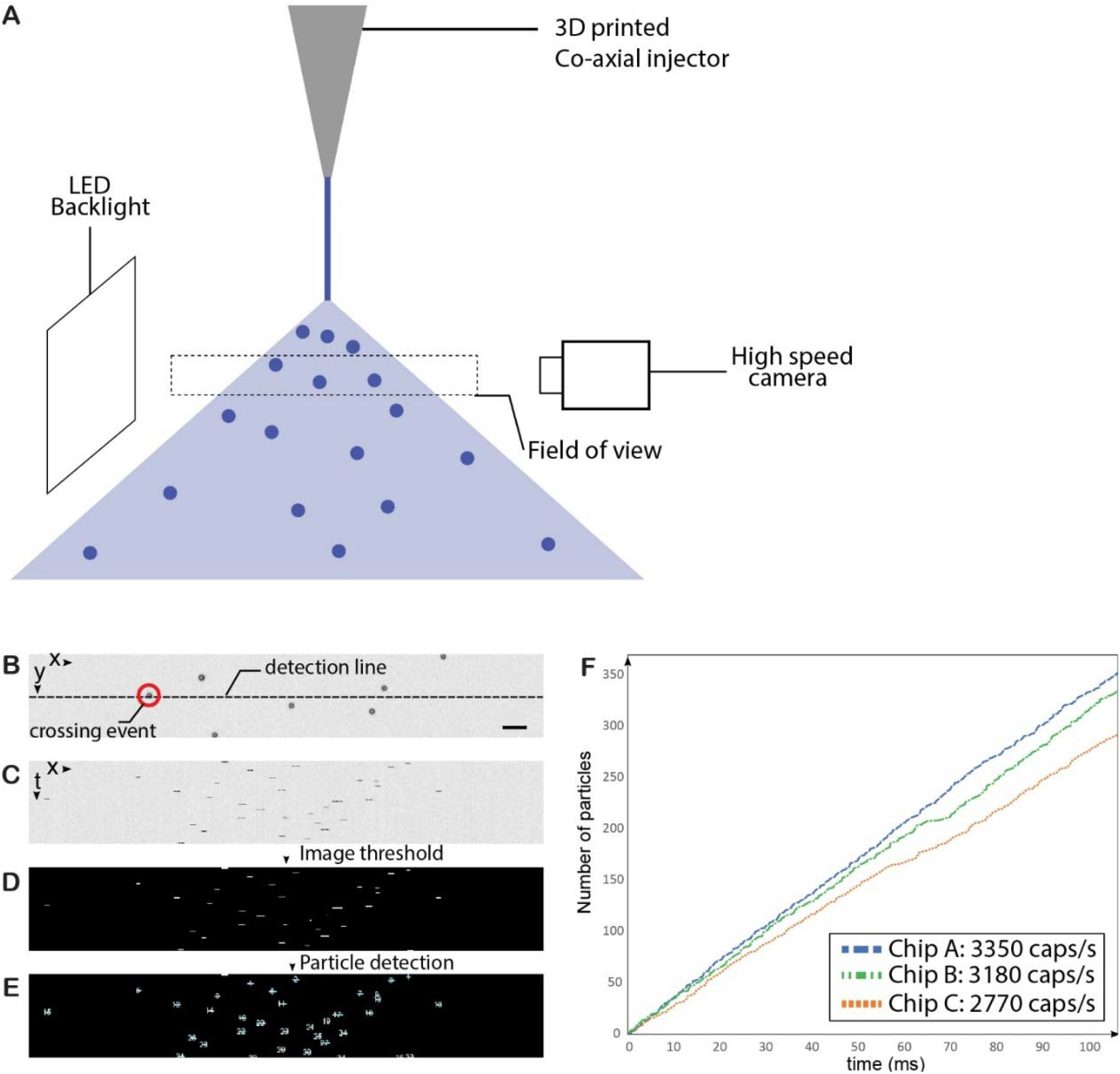
Optical measurement of the capsule production rate. (**A**) Sketch of the experimental setup. The capsules exiting the nozzle are illuminated with a LED backlight (PHLOX 200mx200mm) and imaged using a high-speed camera (HSC Phantom VEO1310L) at a frame rate of 10,000 fps and a spatial resolution is 30px/mm. (**B**) Snapshot showing a typical shadow image of the capsule spray. Scale bar is 1mm. The algorithm increments the account of counts capsules every time one of them crosses the detection line (dashed-white in (b)), and stores the time of the event. Details of the detection algorithm are shown proposed in (c-e). (**C**) Space-time representation of the crossing events, evidenced by stacking the intensity variations of the detection line over time. Each crossing event corresponds to a black spot in the image. (**D**) Intensity threshold of (c). Elementary binary operations are performed to ensure that each crossing event corresponds to a single white spot. Particle analysis (using ImageJ plugin). Each white spot is counted independently, and its coordinates are stored by the algorithm, providing details on the time and location of the crossing event. Example of typical production curves for three different spraying nozzles (chips A, B and C). The corresponding production rates are included in the legend. The values are usually estimated with longer datasets (recordings >500 ms).

**Fig. S2.**
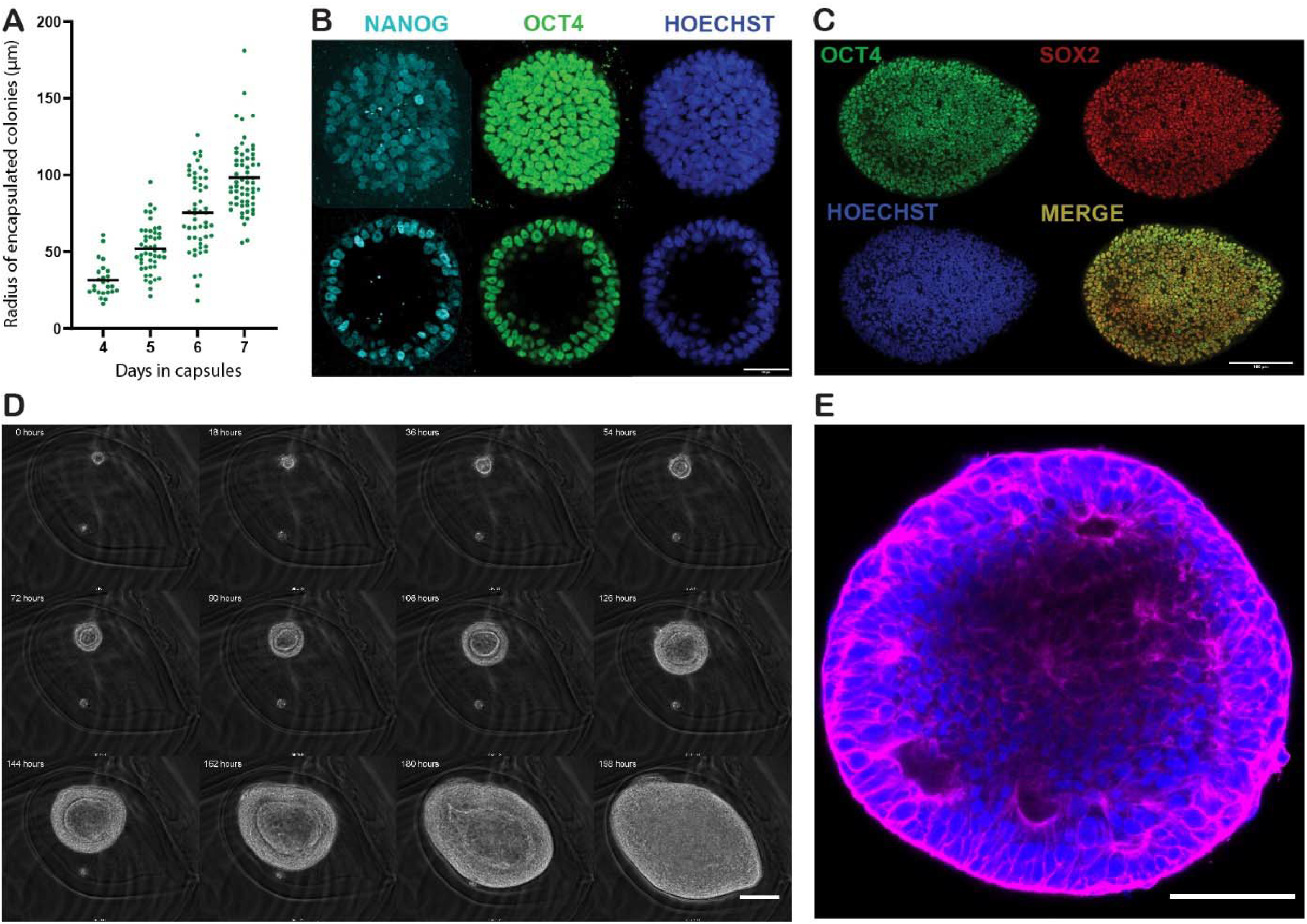
Phenotype of encapsulated 3D hPSC colonies. (**A**) Graph of hPSC cyst radius as a function of time. Radii were measured by monitoring n=25, 47, 52, 61 capsules at days 4, 5, 6 and 7 respectively. (**B**) 3D hPSC colony co-stained for stemness markers NANOG (cyan) and OCT4 (green) and with Hoechst (blue). Projection of maxima (upper panel) and equatorial plane (lower panel) of confocal images. Scale bar is 50µm. (**C**) Confocal equatorial plane of a fully confluent hPSC capsule (t=10 days post-encapsulation) co-stained for OCT4 (green), SOX2 (red), and nuclei (Hoechst, blue) and showing that the lumen has collapsed. Scale bar is 100µm. (**D**) Sequence of phase-contrast microscopy images showing luminogenesis, growth and collapse of an encapsulated epiblast-like colony. The time interval between successive images is 18h. Scale bar=100µm. (**E**) Confocal equatorial plane of a collapsed 3D hPSC colony co-stained for F-actin (Phalloidin, magenta) and nuclei (Hoechst, blue), Scale bar=100µm.

**Fig. S3.**
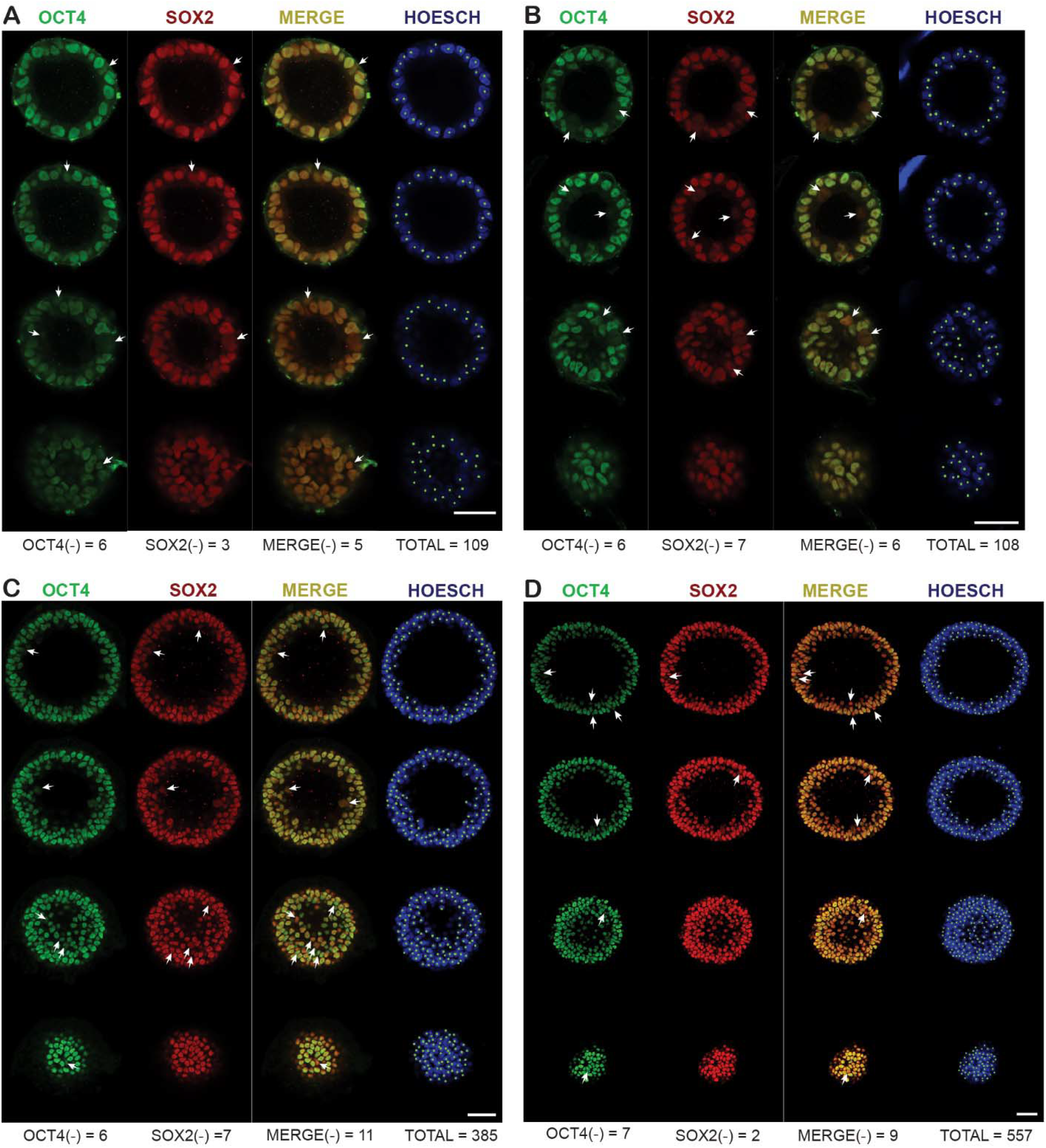
Stemness staining of encapsulated 3D hPSC colonies. Four different representative encapsulated hPSC colonies co-stained for OCT4 (green) SOX2 (red) and nuclei (Hoechst, blue). The 2 colonies in the upper panels (**A**) and (**B**) are collected at day 5. The 2 colonies in the lower panels (**C**) and (**D**) are collected at day7. White arrows and dots illustrate the quantification method used to determine OCT4 (green) and SOX2 (red) co-staining and the percentage of positive cells within the encapsulated cysts shown in Figure 2D of the main text. Scale bars are 50 µm.

**Fig. S4.**
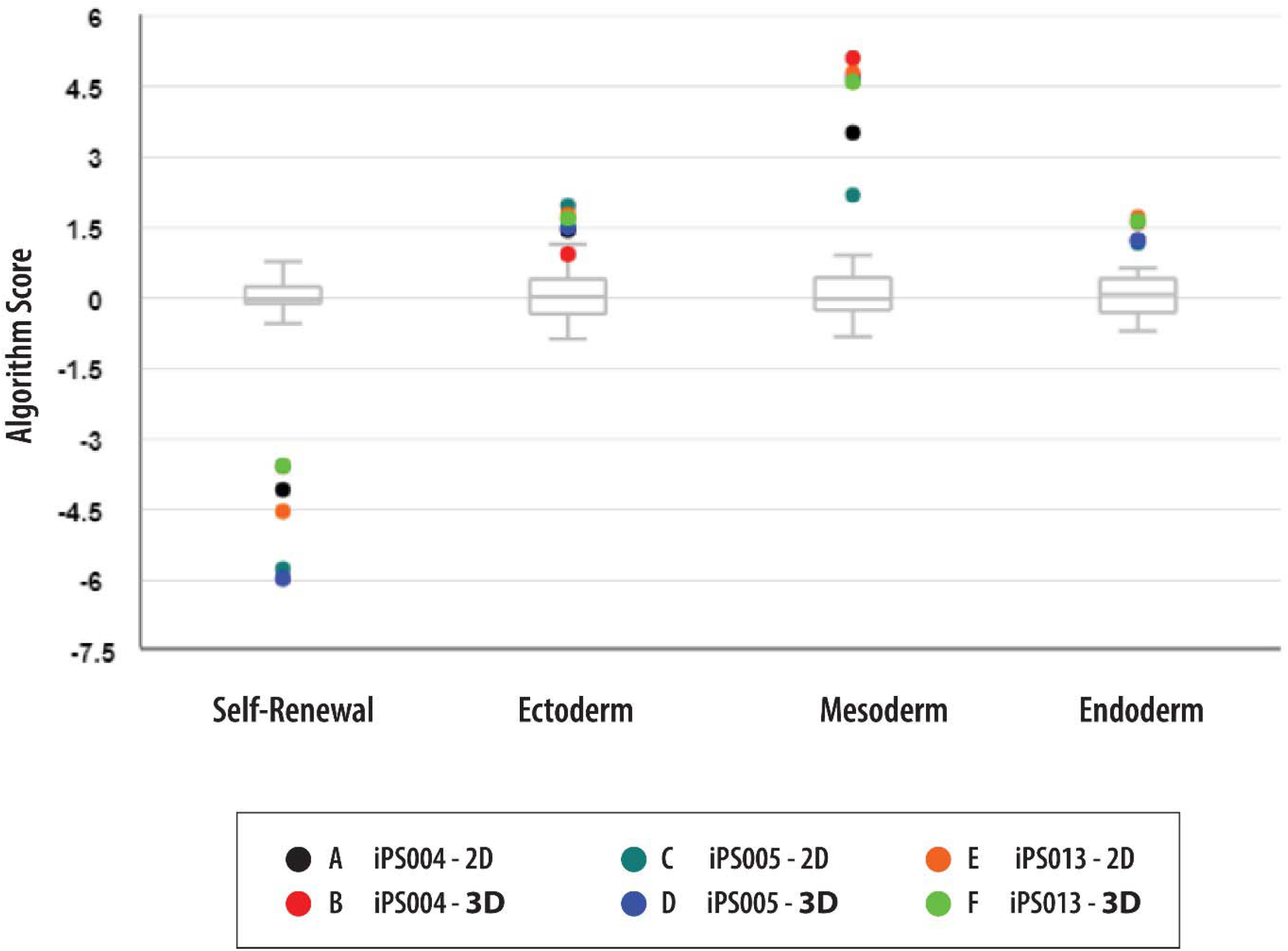
Scorecard quantitative comparison of gene expression profiles from trilineage differentiation assays of 2D and 3D hPSC colonies. 2D cultures and 3D *in capsulo* cultures of hPSC were assessed with trilineage differentiation assay and scorecard qPCR panel. Three iPSC lines were differentiated and analyzed. Algorithm score generated by the manufacturer confirm a consistent decreased of the self-renewal genes compared to the undifferentiated reference. Up-regulation of the 3 germs layers means that both 2D and 3D stem cell colonies can engage in differentiation. The grey box plot indicates the undifferentiated reference.

**Fig. S5.**
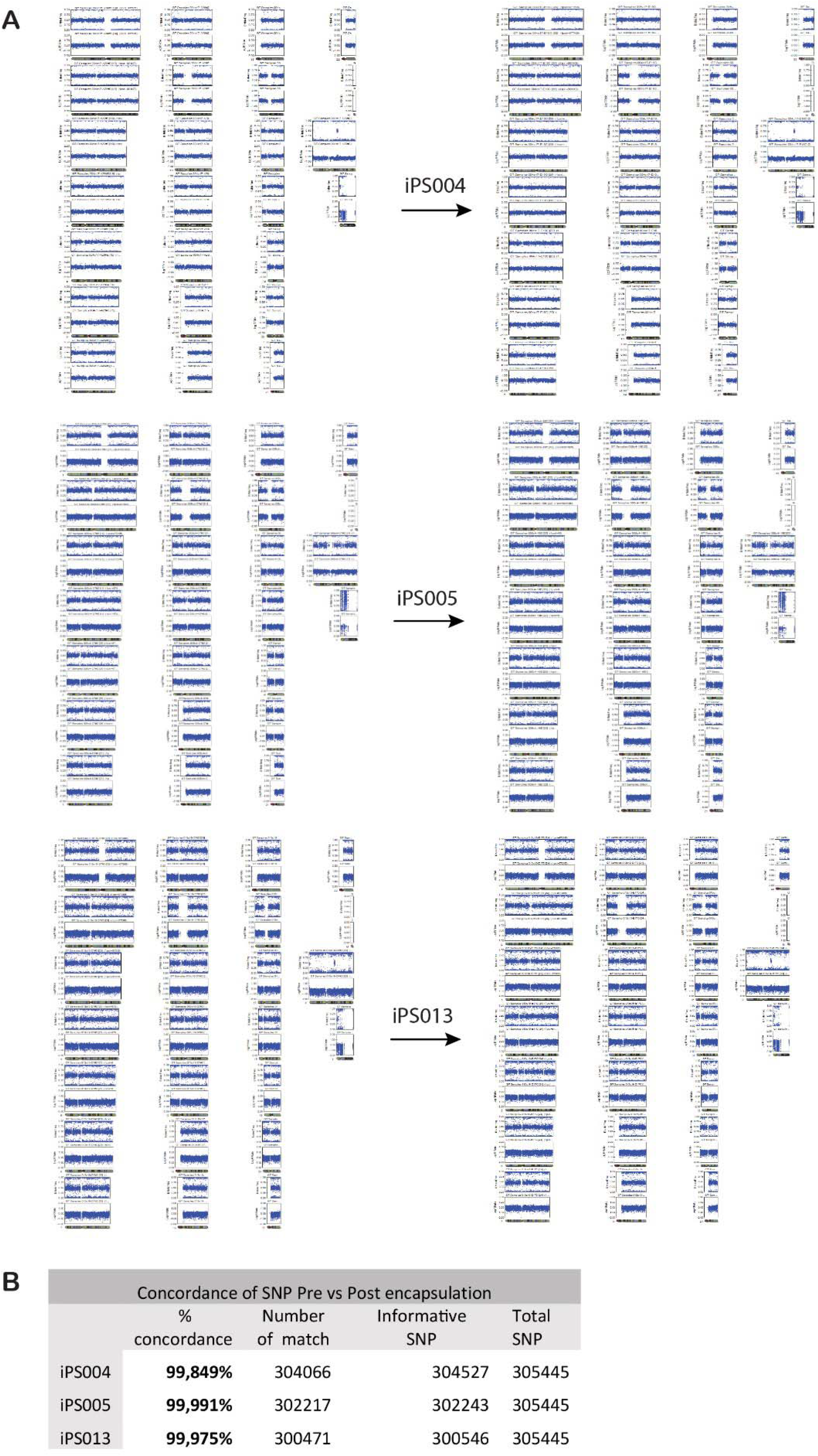
Analysis of high-resolution SNP arrays before and after C-STEM amplification. (**A**) Qualitative analysis of high-resolution SNP arrays before and after one-week of encapsulation: pre-encapsulation (left) and post-encapsulation (right). Absence of duplications and deletions for the 3 distinct iPS cell lines (iPS004 top row, iPS005 middle row, iPS013 lower row). (**B**)Table summarizing the SNP concordance analysis.

**Fig. S6.**
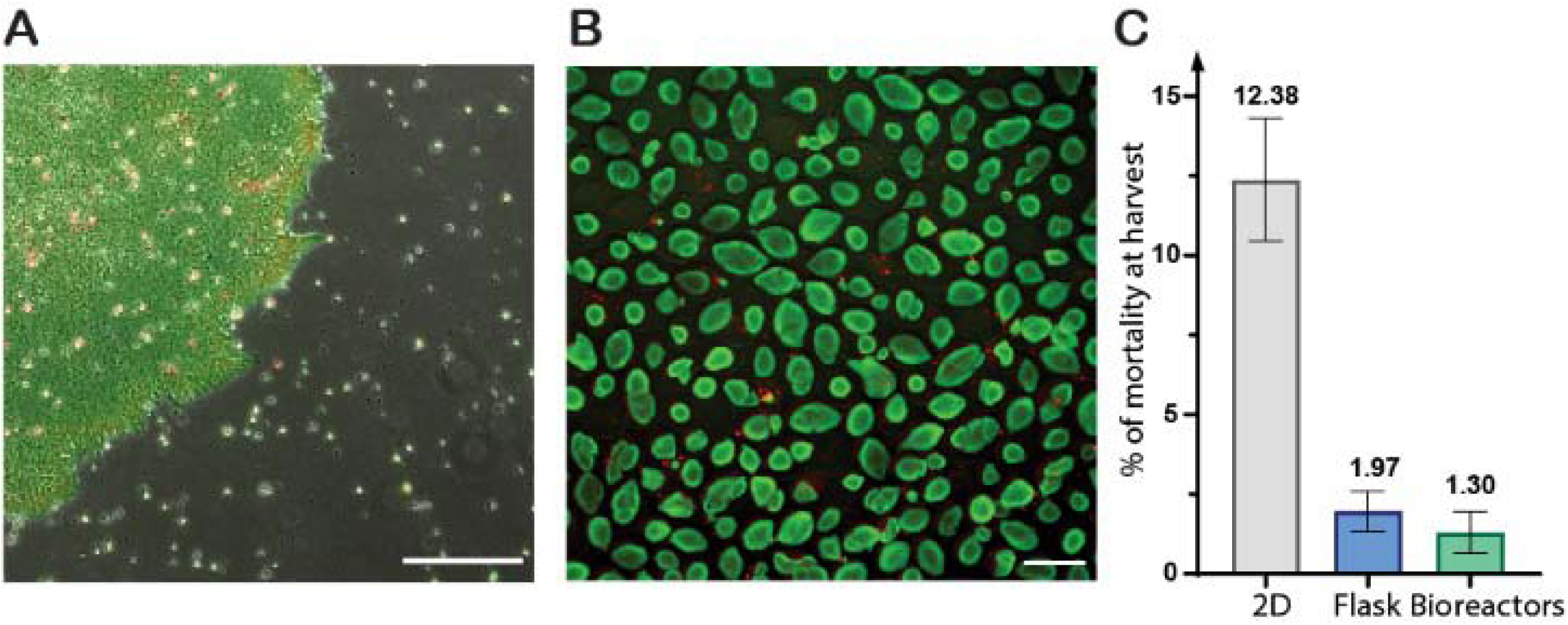
Cell viability in 2D and encapsulated 3D hPSC cultures. (**A**) Fluorescence microscopy image of a 2D stem cell colony stained for live (Calcein, green) and dead (ethidium, red). Pictures were taken before daily media changes to avoid removal of free-floating cells. Scale bar is 500 µm. (**B**) Fluorescence microscopy image of live (green) & dead (red) staining of 3D hPSC colonies at day 7 post encapsulation. Scale bar is 500 µm. (**C**) Percentage of cell mortality at harvest for 2D culture and. encapsulated 3D hPSC colonies. Stem cells cultivated in 2D were harvested at day 4, 5 or 6 before reaching confluency. Stem cells cultivated for 6 to 7 days in capsules were harvested and dissociated after capsule dissolution. Viability was assessed with Nucleo counter NC3000. The ‘count and viability assay’ use the cell stain Acridine Orange for cell detection, and the nucleic acid stain DAPI for detecting non-viable cells.

**Fig. S7.**
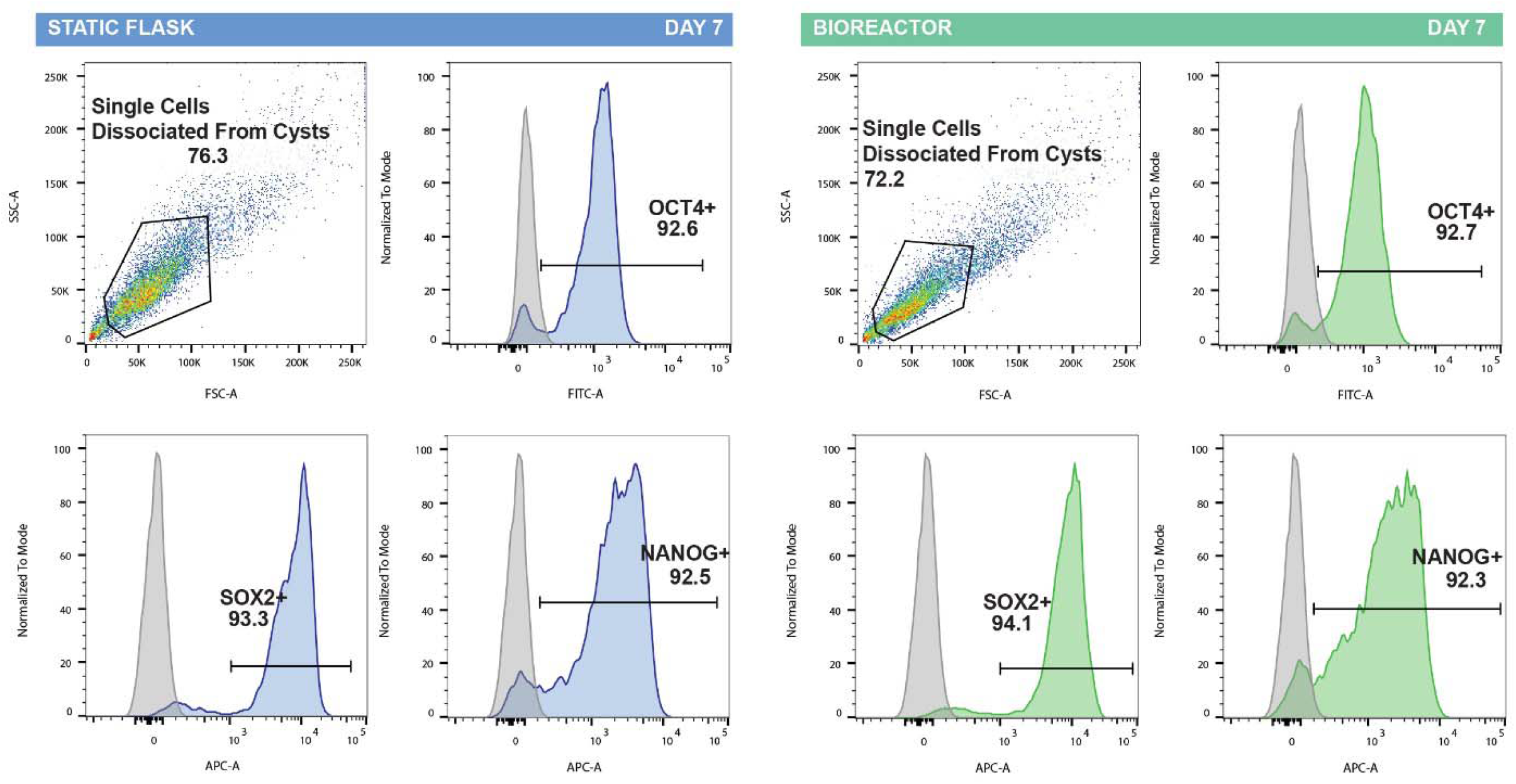
Encapsulated epiblast-like colonies resilience to hydrodynamic damages. From one encapsulation batch, 2 suspension cultures were seeded in parallel: capsules were cultivated either in static suspension (flasks) or in constantly agitated suspension (500ml stirred tank bioreactor). after 7 dyas of suspension culture, capsules were collected, dissolved; 3D stem cell colonies were dissociated, fixed and stained for stem cell markers OCT4, SOX2 and NANOG. Fig. S7 shows flow cytometry dot-plots of the two different culture conditions (static flask versus stirred suspension in a bioreactor)

**Fig. S8.**
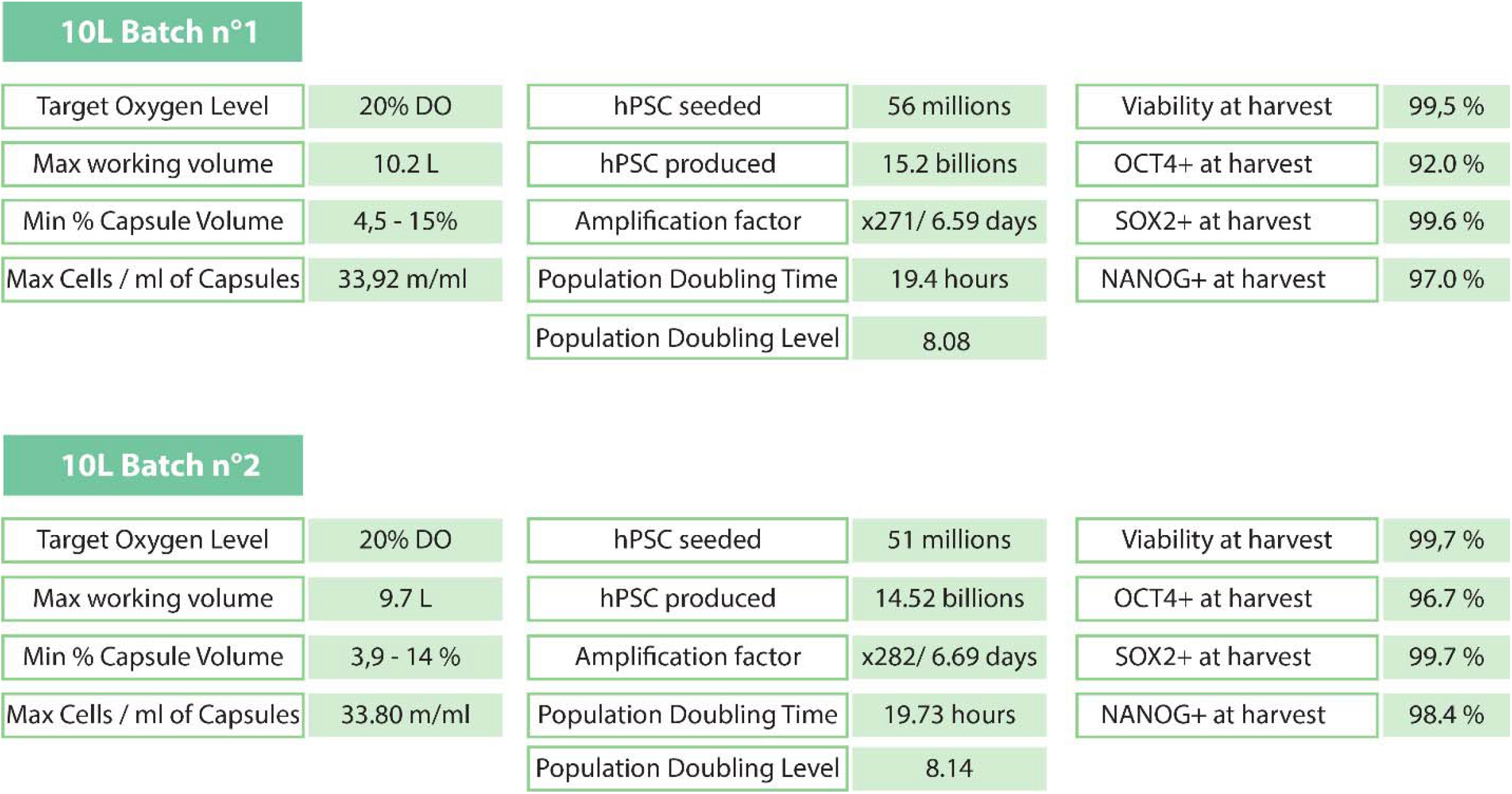
Key parameters and results of C-STEM scale-up in 10 liter bioreactor. Encapsulation of hiPSCs and suspension culture was performed for 2 separated runs. Target oxygen level of 20% Dissolved Oxygen is described as an hypoxic condition (conversely to bioreactors with 100% dissolved oxygen level that are described as normoxic). A fed batch feeding strategy was applied resulting in increasing working volume and decreasing capsule concentration. The cell density can be expressed in millions of cells per milliliter of capsules. The volume of capsule harvested being measured in graduated glass cylinder and reported to the cell quantity. Amplification factor (AF), and Population doubling time (PDT) are defined following main text description. Population doubling level (PDL) is defined as PDL(Dt)= ln(AF) / ln(2). Viability was assessed by nucleocounter NC-3000 (Chemometech). Decapsulated and dissociated cells were fixed and stained for OCT4, SOX2 and NANOG and analyzed by flowcytometry, BD Accuri C6 plus (Table S1).

**Fig. S9.**
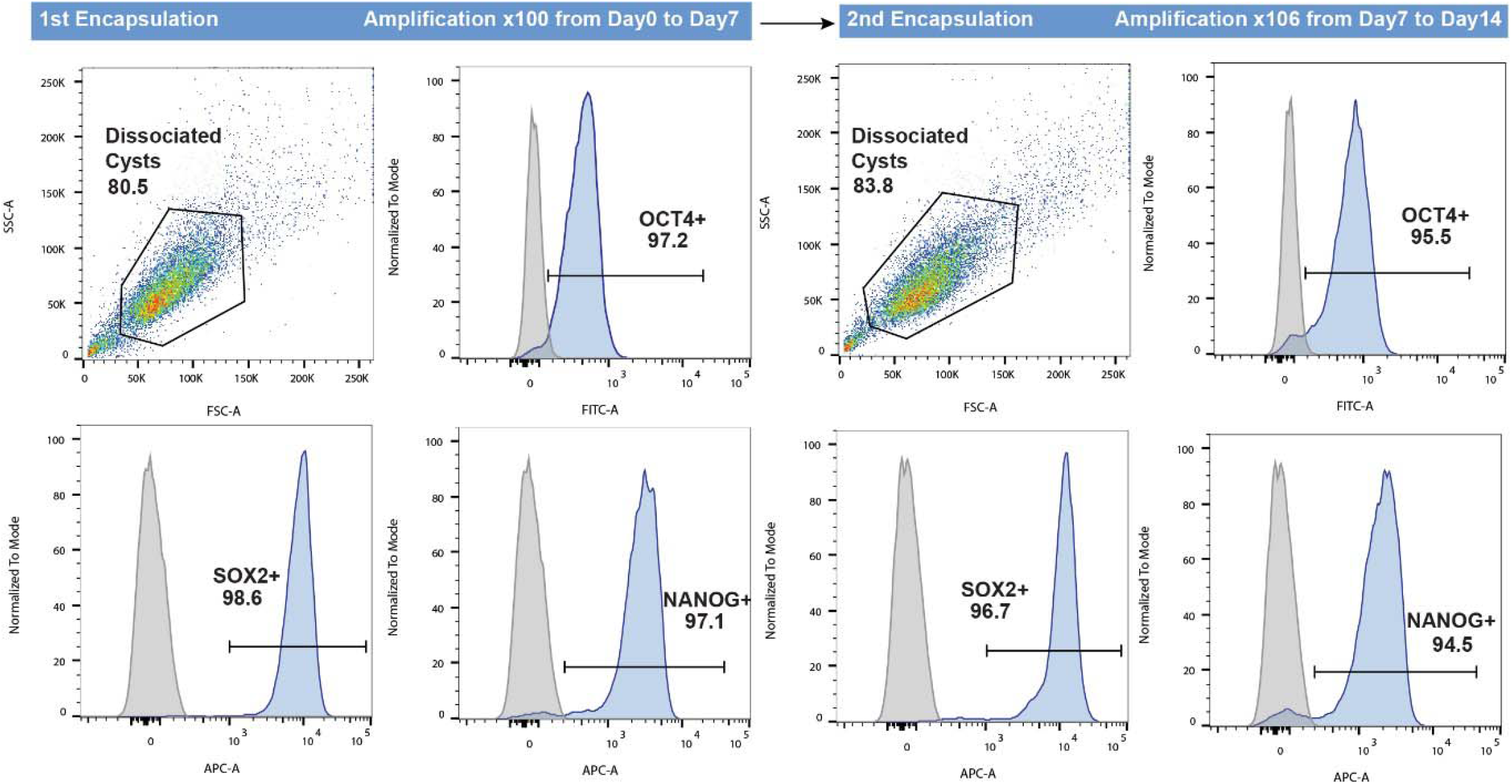
Stemness maintenance assessment of hiPSCs through 2 consecutive encapsulations. After a first encapsulation of hPSCs and 7 days of suspension culture, capsules were collected and dissolved, and 3D colonies were dissociated, resuspended and used for another round of encapsulation/suspension culture. At day 7 and 14 cells were dissociated, fixed, and stained for the stemness markers OCT4, SOX2, and NANOG. Individual histograms from flow cytometry show stable phenotypes.

**MOVIE S1. Formation and collection of capsules in calcium bath.**

Video taken with a high speed camera showing how the train of droplets splays during the encapsulation process under electric field. Acquisition was performed at a frame rate of 10.000 fps. Scale bar is 1mm.

**MOVIE S2. 3D rendering of an equatorially sectioned 3D hPSC colony.**

The image shows nuclear localization of OCT4 (green) and phalloidin (gray). The diameter of the cyst is 110 um.

**MOVIE S3. Time-lapse of an encapsulated 3D hPSC colony : between encapsulation and harvest** Phase contrast sequence (using Biostation Nikon IM) of a growing hPSC colony inside a capsule. Video starts at day 1 after encapsulation. Scale bar is 100 µm.

**MOVIE S4. Time-lapse of an encapsulated 3D hPSC colony : between encapsulation and lumen collapse**

Phase contrast sequence (using Biostation Nikon IM) of a growing hPSC colony inside a capsule. Video starts at day 1 after encapsulation. prolonged until full collapse of the lumen. Scale bar is 100 µm.

**MOVIE S5. Encapsulated 3D hPSCs colonies cultured in a 10 liter stirred-tank bioreactor.** Video taken with a Digital Single-Lens Reflex camera shows the flow of hPSC-laden capsules induced by the motion of the bioreactor’s impeller.

**Table S1.** Antibody list. List of antibodies used for flow cytometry, trilineage assay and immune fluorescence microscopy.

| Antibodies used for flow cytometry |  |
| --- | --- |
| Nanog Hu – Alexa Fluor647 Clone N3-355 | BD (Ref. 561300) |
| OCT3/4 Hu, Ms – Alexa Fluor 488 – clone 40/oct3 | BD (Ref. 560253) |
| SOX2 Hu Alexa Fluor 647 – Clone O30-678 | BD (Ref. 562139) |
| Mouse igG1k – Alexa Fluor 488 – clone MOPC 21 | BD (Ref. 557721) |
| Mouse igG1k, Alexa Fluor 647 clone MOPC 21 | BD (Ref. 557732) |

| Antibodies used for trilineage assay |  |
| --- | --- |
| Monoclonal anti-Tubuline beta 3 (Mouse) 1:1000 | Biolegend Cat#801202 |
| Monoclonal anti actin, alpha-smooth muscle (Mouse) – FITC 1:300 | Sigma-Aldrich Cat#F3777 |
| Monoclonal anti-Alpha foeto protein (Mouse) 1:60 | R&D system Cat#MAB1369 |
| Goat anti-Mouse alexa Fluor plus 555 1:1000 | Thermo fisher Scientific Cat#A32732 |

| Antibodies used for immunostaining and microscopy |  |
| --- | --- |
| OCT 3/4 mouse IgG2b (1/200) | Santa Cruz Biotechnology sc-5279 E1818 |
| SOX2 Rabbit Polyclonal (1/1000) | Sigma Aldrich AB5603 3153252 |
| Secondary Donkey anti-mouse IgG H+L Alexa Fluor Plus 488 1/500 | Thermo/Invitrogen A32766 TF271737A |
| Secondary donkey anti-rabbit IgG H+L Alexa Fluor Plus 555 1/500 | Thermo/Invitrogen A32794 TH271030 |
| Phalloidin Alexa Fluor 647 at 1:500 | Thermo A22287 2015553 |
| Hoechst 33342 at 1:1000 |  |

