## Supplementary figures and images for "C-STEM: ENGINEERING NICHE-LIKE MICRO-COMPARTMENTS FOR OPTIMAL AND SCALE-INDEPENDENT EXPANSION OF HUMAN PLURIPOTENT STEM CELLS IN BIOREACTORS"

### Fig. S1. Optical measurement of the capsule production rate.

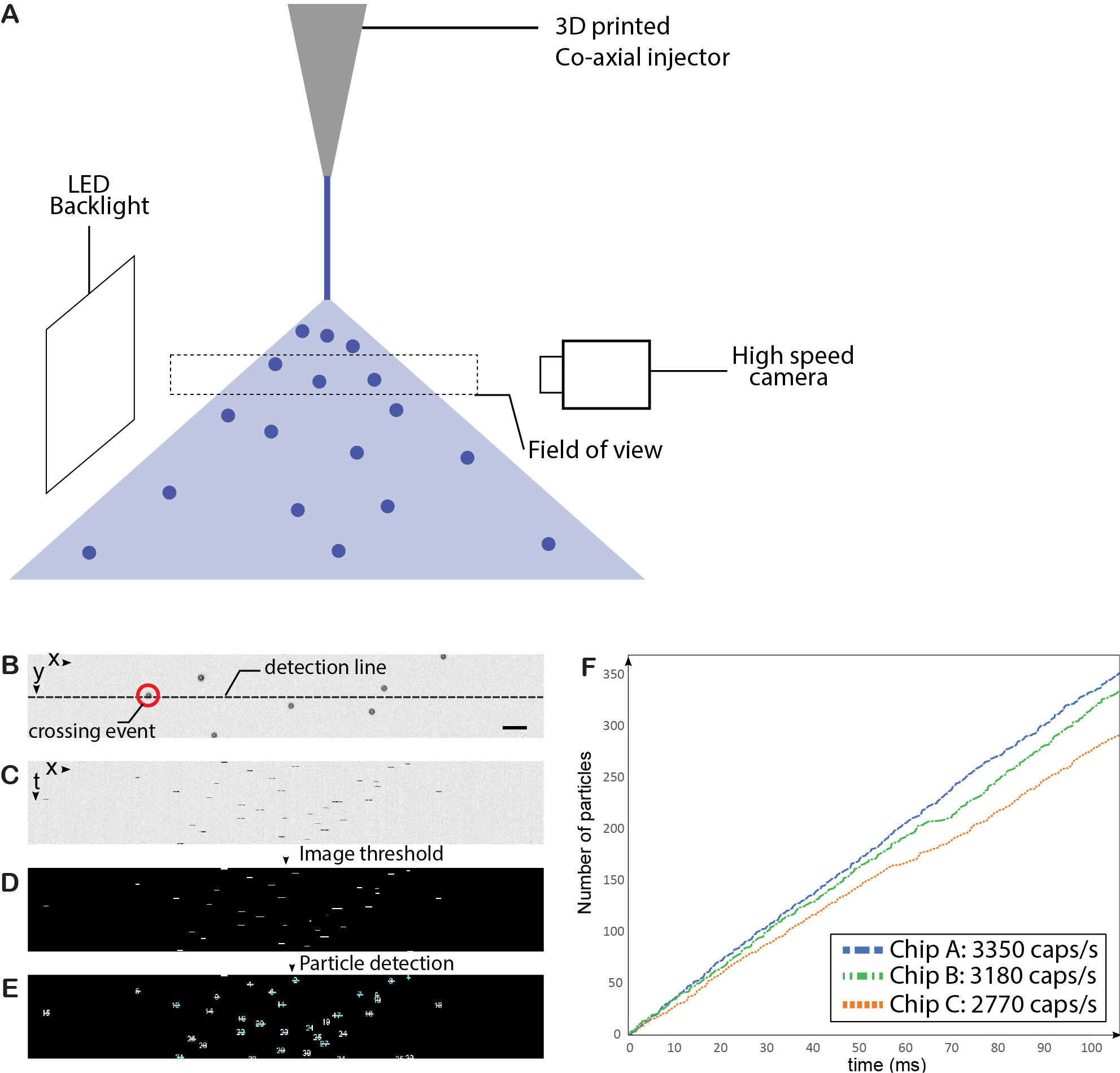

### Fig. S2. Phenotype of encapsulated 3D hPSC colonies

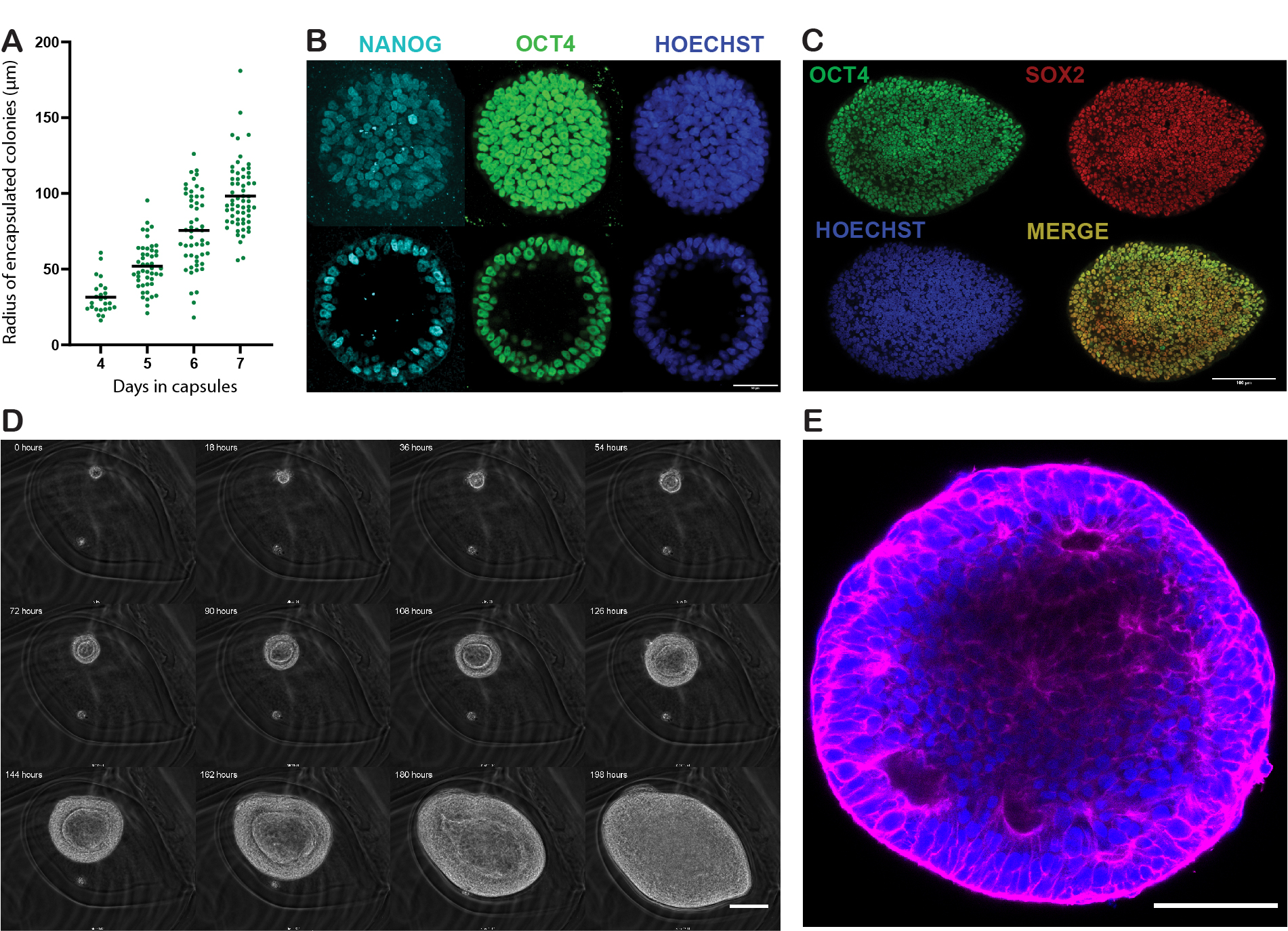

### Fig. S3. Stemness staining of encapsulated 3D hPSC colonies

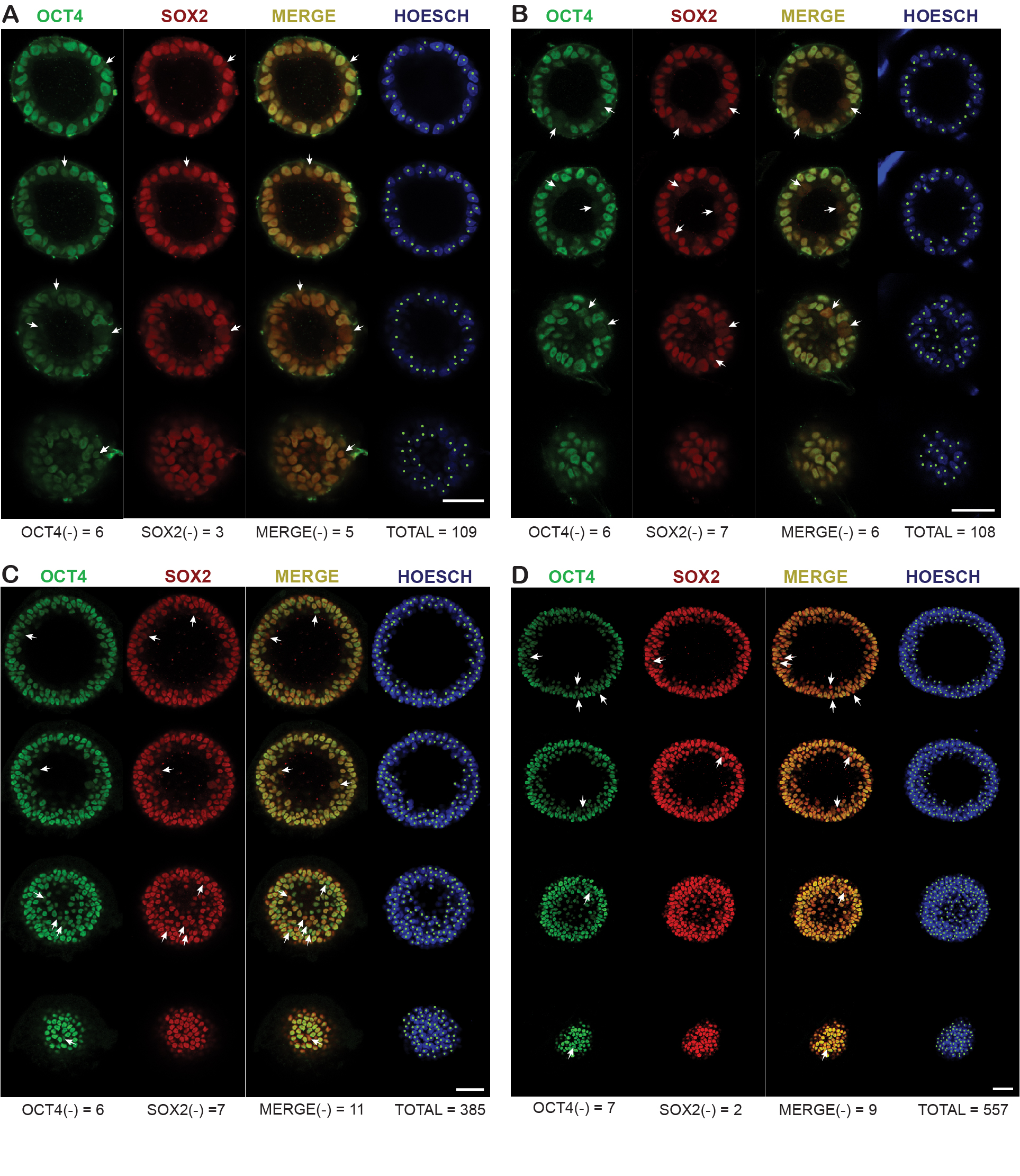

### Fig. S4. Scorecard quantitative comparison of gene expression profiles from trilineage differentiation assays of 2D and 3D hPSC colonies

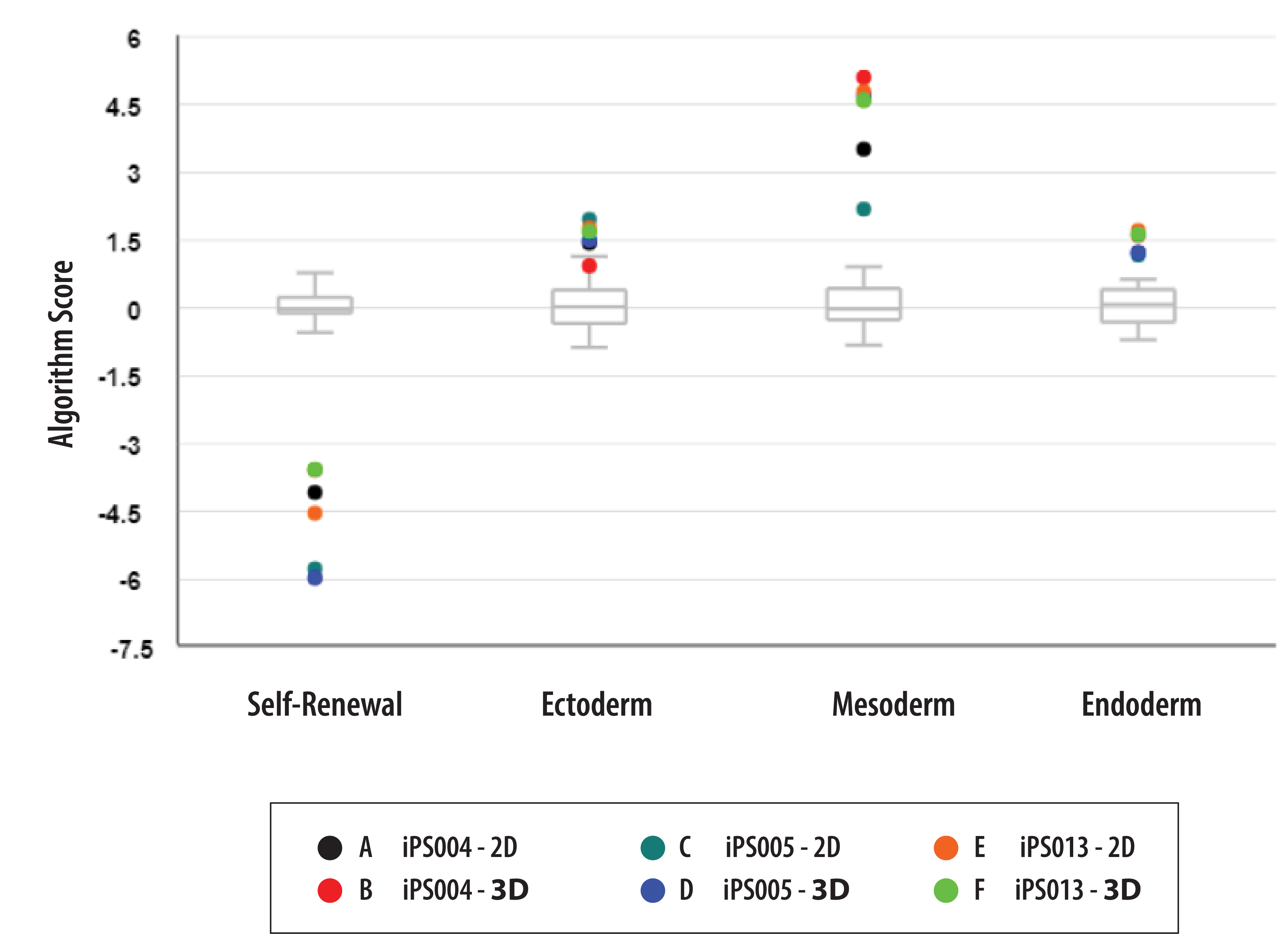

### Fig. S5. Analysis of high-resolution SNP arrays before and after C-STEM amplification

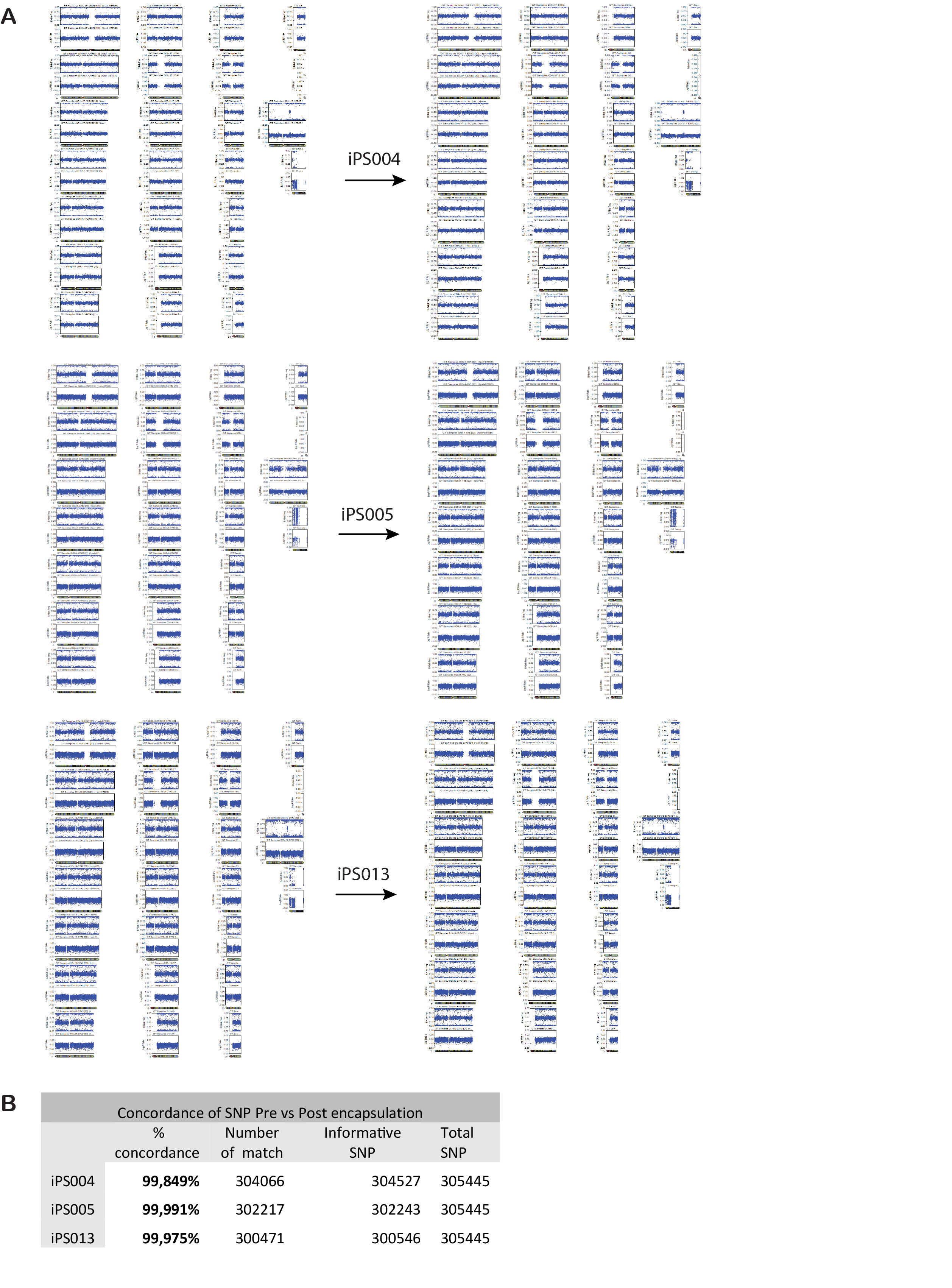

### Fig. S6. Cell viability in 2D and encapsulated 3D hPSC cultures

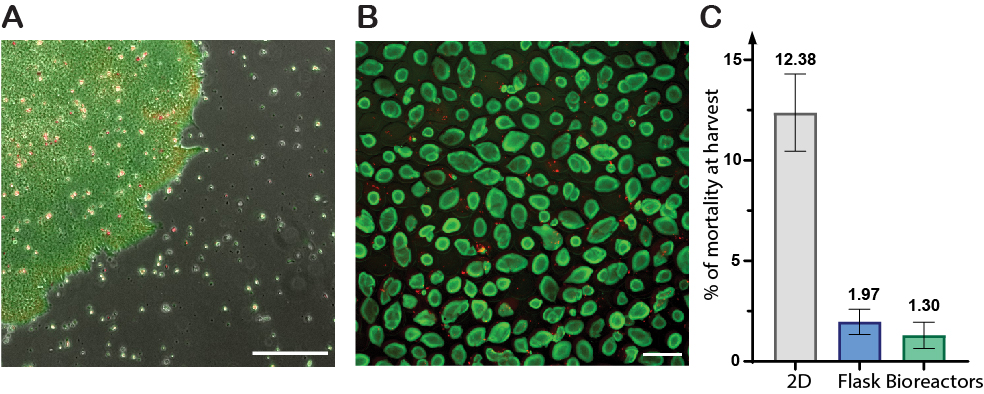

### Fig. S7. Encapsulated epiblast-like colonies resilience to hydrodynamic damages

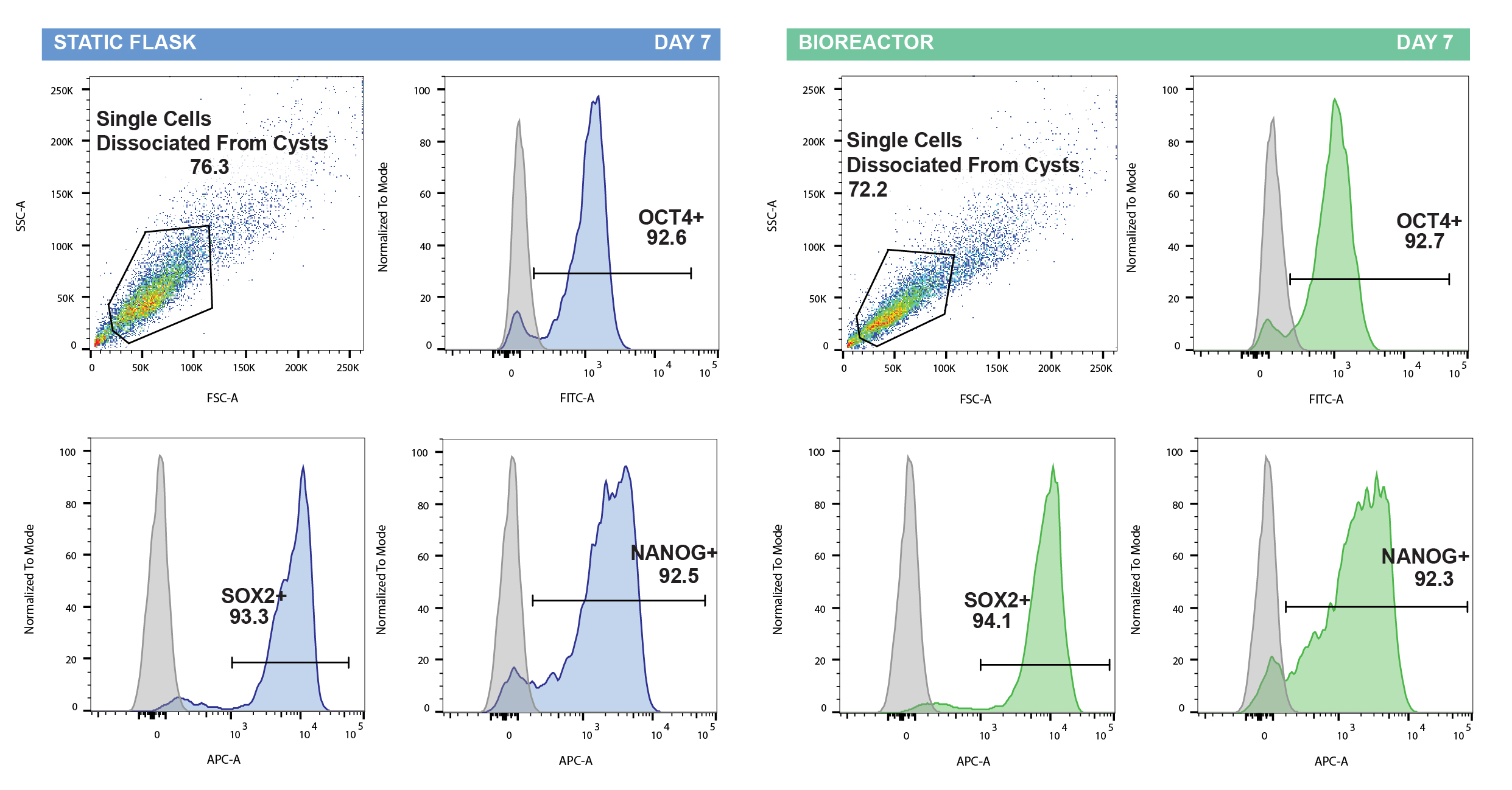

### Fig. S8. Key parameters and results of C-STEM scale-up in 10 liter bioreactor

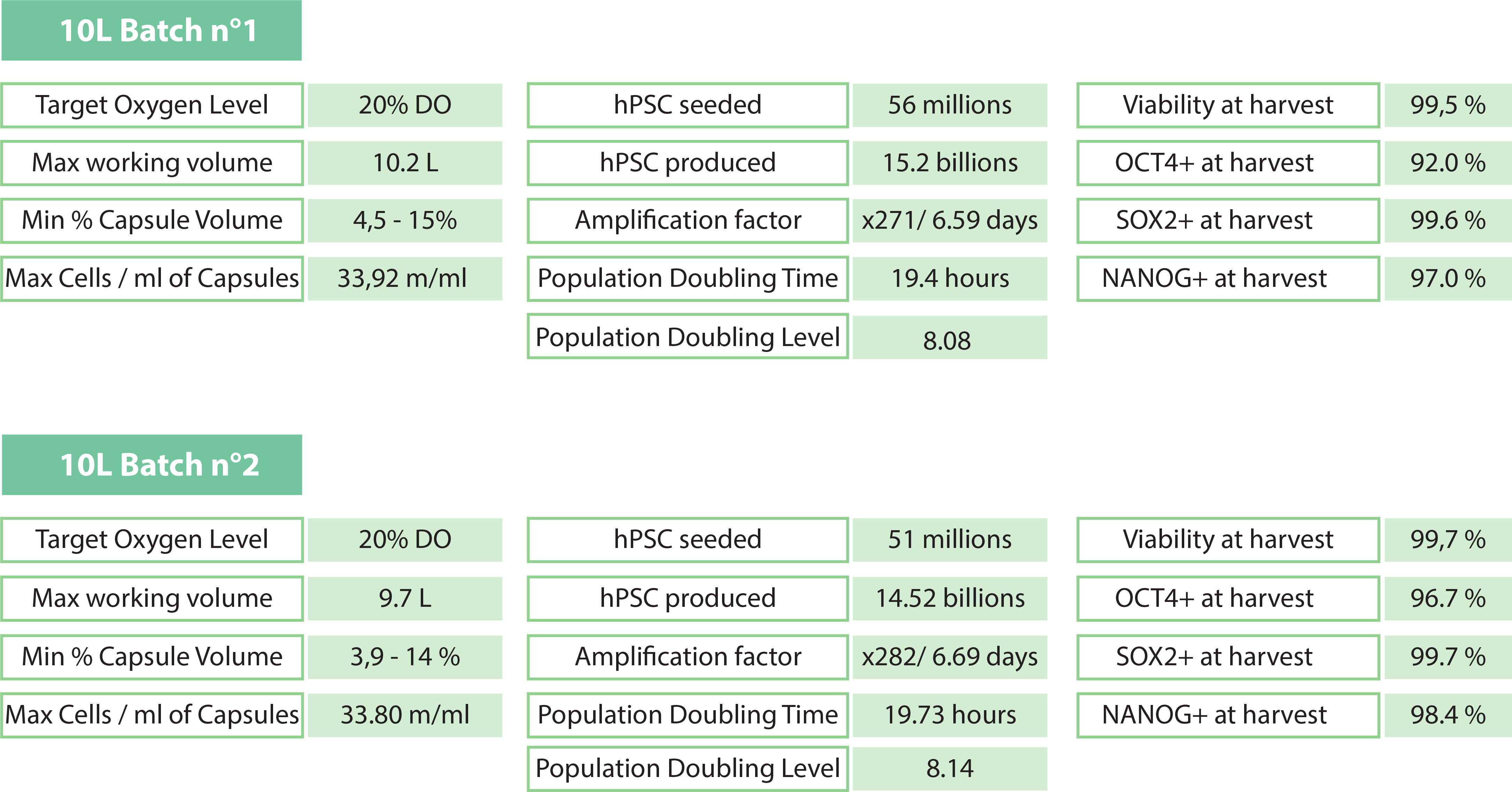

### Fig. S9. Stemness maintenance assessment of hiPSCs through 2 consecutive encapsulations

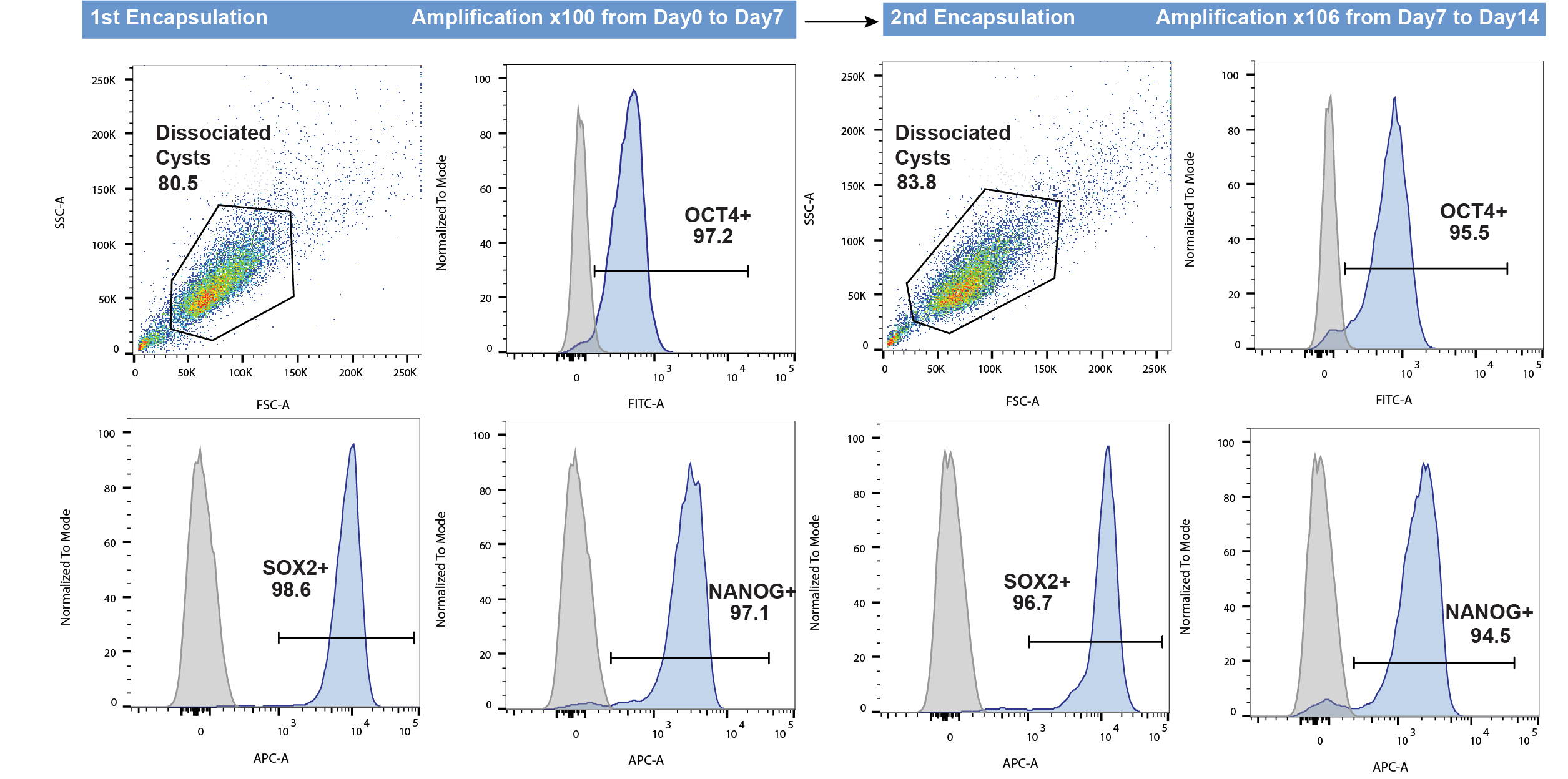
